# CARP Protects Against Doxorubicin-induced Cardiotoxicity through NRF1-driven Mitochondrial Homeostasis

**DOI:** 10.1101/2025.11.06.683842

**Authors:** Dan Zhong, Xusan Xu, Taoshan Feng, Wensen Zhang, Fei Luo, Junhao Guo, Zhengqiang Luo, Riling Chen, Yajun Wang, Guoda Ma

## Abstract

**Aims:** Doxorubicin (DOX), a highly effective anthracycline chemotherapeutic agent, is limited by its dose-dependent cardiotoxicity. This study investigates the cardioprotective mechanisms of cardiac adriamycin responsive protein (CARP) and its underlying mechanisms in DOX-induced cardiotoxicity (DIC).

**Methods and results:** Cardiac-specific CARP transgenic and wild-type mice were subjected to a DIC model. Cardiac function was assessed via echocardiography, histopathology, and transmission electron microscopy (TEM). *In vitro*, DOX-treated cardiomyocytes overexpressing CARP were analyzed for oxidative stress (ROS levels), mitochondrial function (mtDNA copy number etc.), and mitochondrial-related proteins (Western blot). CARP-NRF1 interaction was validated by co-immunoprecipitation (Co-IP), and NRF1 siRNA knockdown was performed to assess CARP’s role in mitochondrial homeostasis. CARP overexpression markedly alleviated DOX-induced cardiac dysfunction and mitochondrial damage, restoring mitochondrial dynamics and mitophagy. Mechanistically, CARP directly interacted with NRF1, and NRF1 knockdown ameliorated CARP-mediated cardioprotection in DIC.

**Conclusion:** CARP safeguards against DIC by maintaining mitochondrial homeostasis via NRF1 signaling, positioning it as a promising therapeutic target for DOX-induced cardiomyopathy.

## 1 Introduction

Doxorubicin (DOX), an anthracycline antibiotic first isolated in 1967 from *Streptomyces peucetiusmutant* strains ^[1]^, remains a potent chemotherapeutic agent for both solid and hematologic malignancies across age groups due to its broad-spectrum antitumor efficacy ^[2, 3]^. However, its clinical utility is significantly limited by dose-dependent cardiotoxicity ^[4]^. While current strategies—including optimized dosing regimens, iron chelation therapy, and conventional heart failure medications—provide partial protection against DOX-induced cardiotoxicity (DIC) ^[5]^, they fail to address the fundamental pathological mechanisms. This critical gap underscores the urgent need to elucidate the molecular pathogenesis of DIC to develop targeted therapies.

Emerging research has established mitochondrial dysfunction as the central pathological mechanism in DIC ^[6, 7]^. Cardiomyocytes exhibit susceptibility to oxidative damage due to their high mitochondrial content and relatively limited antioxidant capacity ^[8, 9]^. This vulnerability is dramatically amplified by DOX’s unique pharmacokinetics: the drug accumulates in mitochondria at concentrations up to 100-fold higher than plasma levels ^[7]^. Within mitochondria, DOX binds to cardiolipin, disrupts electron transport chain function, and initiates a cascade of metabolic dysfunction and excessive reactive oxygen species (ROS) production, ultimately leading to irreversible myocardial injury ^[10, 11]^.

The maintenance of mitochondrial homeostasis relies on precisely coordinated quality control mechanisms, including dynamic fission/fusion processes, selective mitophagy, and biogenesis ^[12, 13]^. DOX exposure disrupts this delicate balance, inducing pathological mitochondrial fission and excessive mitophagy that promote cardiomyocyte death and cardiac dysfunction ^[14, 15]^. Therapeutic interventions targeting these processes show considerable promise: genetic ablation of dynamin-related protein 1 (DRP1), the master regulator of mitochondrial fission or pharmacological inhibition of mitochondrial fission (e.g., using mitochondrial division Inhibitor 1, Mdivi-1) improves mitochondrial function and mitigates DIC ^[16, 17]^. Similarly, knockdown of parkin RBR E3 ubiquitin protein ligase (Parkin) attenuates hyperactivated mitophagy and consequently reduces DOX-induced cardiomyocyte death ^[18]^. Intriguingly, while mitophagy eliminates damaged organelles, it also serves as a crucial mechanism for mitochondrial turnover and renewal ^[19, 20]^. Complementary studies demonstrate that enhancing mitochondrial biogenesis effectively counteracts DOX-induced myocardial injury ^[21, 22]^. Notably, recent advances in mitochondrial transplantation have shown remarkable potential in ameliorating DOX-induced cardiac damage ^[23–25]^. Together, these findings position mitochondrial quality control modulation as a highly promising therapeutic strategy for DIC ^[7, 26]^.

Cardiac adriamycin-responsive protein (CARP), alternatively termed cardiac ankyrin repeat protein (CARP) or ankyrin repeat domain 1 (ANKRD1), is a heart-specific multifunctional protein with dual physiological functions. Nuclear CARP functions as a transcriptional cofactor, interacting with regulators including nuclear factor-kappa B (NF-κB) and Y-box binding protein 1 (YB-1) to modulate gene expression ^[27–29]^, while at the sarcomeric I-band, it maintains myocardial structural integrity and contractility ^[30]^. Notably, CARP expression is rapidly downregulated in DOX-treated cardiomyocytes ^[31, 32]^ and our clinical observations demonstrate reduced serum CARP levels in acute lymphoblastic leukemia patients with anthracycline-induced cardiac adverse reaction ^[33]^, implicating its potential role in DIC pathogenesis. Current evidence suggests that CARP participates in the chronic activation of G protein-coupled estrogen receptor 1 (GPER1), mediating antioxidant stress responses and mitochondrial protection in cardiomyocytes ^[34]^. Mechanistically, CARP upregulates the anti-apoptotic protein B-cell Lymphoma 2 (Bcl-2) to counter hypoxia-induced apoptosis ^[35]^. These findings collectively indicate that CARP-mediated cardioprotection may involve mitochondrial regulation, though its specific role in maintaining mitochondrial homeostasis during DIC requires further elucidation.

Mass spectrometry-based screening initially identified a potential interaction between CARP and nuclear factor erythroid 2-related factor 1 (NRF1) ^[36]^, a master regulator of mitochondrial biogenesis ^[37]^. NRF1 coordinates mitochondrial homeostasis through two principal mechanisms: (1) regulating the transcriptional network of nuclear-encoded mitochondrial proteins ^[38]^, and (2) stabilizing mitochondrial DNA copy number while optimizing synthesis of mitochondrial DNA (mtDNA) -encoded proteins ^[39]^. Beyond biogenesis, NRF1 exhibits multifaceted functionality, including antioxidative stress protection ^[40, 41]^, transcriptional regulation of mitophagy ^[42, 43]^ and modulation of fission dynamics ^[44]^.

This study is designed to investigate how CARP-mediated activation of the NRF1 pathway preserves mitochondrial homeostasis, with the goal of validating CARP as a novel therapeutic target for DIC intervention.

## 2 Materials and Method

### 2.1 Ethics statement

All animal experiments were performed in strict accordance with Directive 2010/63/EU of the European Parliament and were approved by the Institutional Animal Care and Use Committee of Guangdong Medical University (Approval ID: GDY1902149). All efforts were made to minimize animal suffering and to reduce the number of animals used . At the study endpoint, all animals were humanely anaesthetized by isoflurane inhalation 2% and then euthanized by cervical dislocation.

### 2.2 Animals and treatment

Male C57BL/6J mice (6–8 weeks old, 20 ± 2 g) were obtained from the Guangdong Medical Laboratory Animal Center (Foshan, China). CARP transgenic mice (CARP-Tg) generated under the control of the α-myosin heavy chain (α-MHC) promoter, were procured from Cyagen Biosciences (Guangzhou, China). Genotyping was conducted via tail DNA extraction followed by PCR amplification.

For Experiment 1: male C57BL/6J mice were randomly divided into two groups (n = 4/group): normal saline (NS) group and doxorubicin (DOX) group. Mice in the DOX group received weekly tail vein injections of DOX (5 mg/kg, Selleck, USA) at weeks 0, 1, 2, and 3, resulting in a cumulative dose of 20 mg/kg over the treatment period, whereas the NS group received equivalent volumes of sterile saline via the same injection protocol ^[45]^. In Experiment 2: male C57BL/6J mice and CARP-Tg mice were randomly assigned to four groups (n = 6/group): wide type (WT)+NS, CARP-Tg, WT+DOX, and CARP-Tg+DOX.

The injection method and doses of doxorubicin and normal saline were the same as those described earlier.

### 2.3 Echocardiography assessment

Cardiac function was evaluated by echocardiography two weeks following the final DOX administration. Under anesthesia (2% isoflurane), mice underwent ultrasound examination using a high-resolution small-animal ultrasound system (VisualSonics, USA). Standard parasternal long-axis views were acquired, and left ventricular functional parameters were derived from M-mode tracings. All functional parameters were derived from the average of 5 consecutive, stable cardiac cycles, with triplicate measurements performed for each animal to ensure reproducibility.

### 2.4 Assessment of myocardial injury

Serum samples were obtained by centrifugation of whole blood at 3,000 × g for 20 min at 4°C. Myocardial injury was evaluated by measuring serum concentrations of creatine kinase (CK), its myocardial-specific isoenzyme (CK-MB), lactate dehydrogenase (LDH), and spartate aminotransferase (AST) using an automated clinical chemistry analyzer.

### 2.5 Transmission electron microscopy (TEM) analysis

Left ventricular tissue samples (1 mm³) were fixed in electron microscopy fixative (Servicebio, China) for 2 h at room temperature and subsequently washed three times with 0.1 M phosphate-buffered saline (PBS, pH 7.4). Post-fixation was carried out in 1% osmium tetroxide for 2 h, followed by additional PBS washes 3 times. The samples were then dehydration through graded ethanol and acetone series and embedded in Pon™ 812 epoxy resin. After overnight incubated at 37°C, the resin-embedded blocks were polymerized at 60°C for 48 h. Ultrathin sections (50 nm) were obtained using an ultramicrotome (Leica, Germany), double-stained with uranyl acetate and lead citrate (15 min each), and examined under a TEM (Hitachi, Japan).

### 2.6 Histopathological analysis

Cardiac tissue samples were fixed in 4% paraformaldehyde (Biosharp, China) followed by graded ethanol dehydration, xylene clearing, and paraffin embedding. Serial sections (5 μm thickness) were prepared and stained with hematoxylin-eosin (H&E) for general morphology, masson’s trichrome for collagen deposition, and wheat germ agglutinin (WGA) for cardiomyocyte membrane visualization. All stained sections were imaged using a fluorescence microscope (Thermo Fisher Scientific, USA) to assess structural and compositional alterations.

### 2.7 Terminal transferase UTP nick end labeling (TUNEL) assay

Paraffin-embedded cardiac tissue sections were sequentially deparaffinized in xylene and rehydrated through a graded ethanol series, followed by three washes with PBS. To facilitate antigen accessibility, sections underwent enzymatic digestion with 20 μg/ml DNase-free proteinase K at 37°C for 20 min. The TUNEL reaction was subsequently performed by incubating sections with terminal deoxynucleotidyl transferase (TdT) enzyme in a light-protected humidified chamber at 37°C for 2 h. Following DNA fragmentation labeling, sections were counterstained with DAPI-containing antifade mounting medium to visualize nuclei. Fluorescence microscopy was employed to detect and quantify apoptotic cells based on TUNEL-positive signals.

### 2.8 Cell culture

The rat cardiomyocyte cell line H9C2, human embryonic kidney cell line HEK293T, and Lenti-293T cells were obtained from the Cell Resource Center of Chinese Academy of Sciences (ATCC, China). All cell lines were cultured in Dulbecco’s Modified Eagle Medium supplemented with 10% fetal bovine serum and 100 U/ml penicillin/streptomycin at 37°C in a humidified 5% CO2 incubator.

### 2.9 Establishment of CARP-overexpressing H9C2 Cell

The rat CARP cDNA was cloned into the lentiviral vector LV-003 (VectorBuilder, China) to generate the pLV-CARP construct, with the empty vector pLV-CON served as a negative control. For lentivirus production, pLV-CARP and pLV-CON were co-transfected with packaging plasmids (psPAX2 and pMD2G) into Lenti-293T cells using Lipofectamine™ 3000 (Thermo Fisher Scientific, USA). The resulting lentiviral particles (LV-CARP and LV-CON) were used to infect H9C2 cardiomyocytes. Stable cell lines were established through 14-day selection with 600 μg/ml geneticin (Beyotime, China). To examine CARP’s role in DIC, cells were divided into four experimental groups: pLV-CON, pLV-CON+0.75 μM DOX (Sigma, USA), pLV-CARP, and pLV-CARP+0.75 μM DOX, with DOX treatment 24 h prior to analysis.

### 2.10 Quantitative real-time PCR (qPCR) analysis

Total RNA was extracted from H9C2 cells and cardiac tissues using TRIzol reagent (TransGen Biotech, China) following the manufacturer’s instructions. For cDNA synthesis, 1 μg of total RNA was reverse transcribed using a reverse transcription kit (NovoProtein, China) with a mixture of random hexamer and oligo (dT) primers. Quantitative PCR amplification was performed using Fast SYBR Green Master Mix (Vazyme, China) on a Quantstudio.5 PCR System (Thermo Fisher Scientific, USA). Expression levels of target genes were normalized to the endogenous reference gene GAPDH. All primer sequences used in this study are provided in Supplementary Table S1.

### 2.11 Western blot analysis

Protein lysates were extracted from cells and cardiac tissues using RIPA buffer (Beyotime, China), and total protein concentrations were determined using a BCA assay kit (Beyotime, China). Equal amounts of protein were separated by SDS-PAGE and subsequently transferred onto PVDF membranes (Millipore, USA). Membranes were blocked with 5% non-fat milk in TBST for 1 h at room temperature, followed by overnight incubation at 4°C with primary antibodies (see Supplementary Table S2 for details). After washing, membranes were probed with appropriate HRP-conjugated secondary antibodies for 45 min at room temperature. Protein bands were detected using enhanced chemiluminescence substrate (Millipore, USA).

### 2.12 Cell viability assay

Following treatment, cell viability was assessed using the CCK-8 assay (GLPBIO, USA). Briefly, 10% CCK-8 reagent was added to each well and incubated at 37°C for 2 h. H9C2 cells were seeded in 96-well plates and treated with DOX. Following treatment, 10% CCK-8 reagent (GLPBIO, USA) was added to each well and incubated at 37°C for 2 h. Absorbance was measured at 450 nm using a microplate reader (BioTek, USA), and relative cell viability was calculated by normalizing to untreated controls.

### 2.13 Mitochondrial function assessment

Mitochondrial membrane potential (MMP) was assessed using the JC-1 assay kit (Beyotime, China) following manufacturer instructions. Briefly, cells were seeded in confocal dishes and exposed to 0.75 μM DOX for 24 h, with 10 μM carbonyl cyanide m-chlorophenyl hydrazone (CCCP) included as a positive control for membrane depolarization. After treatment, cells were loaded with JC-1 probe at 37°C for 20 min under light-protected and washed twice with assay buffer. Fluorescence imaging was performed using laser-scanning confocal microscopy (Leica, Germany).

Cellular ATP levels were determined using a commercial ATP assay kit (Beyotime, China). Following cell lysis, total protein concentration was measured by BCA protein assay. The supernatant was then mixed with ATP working solution and incubated under dark conditions for 30 min. ATP-dependent luminescence was quantified in relative luminescence units (RLU) using a microplate reader, with final ATP concentrations normalized to total protein content.

Total DNA was isolated from cells and cardiac tissues using a DNA extraction kit (Tiangen, China). Relative mtDNA copy number was determined by qPCR using primers specific to the mitochondrial-encoded NADH dehydrogenase subunit 1 (ND1) gene, with nuclear β-actin as the reference gene.The mtDNA abundance was expressed as the ratio of ND1 to β-actin amplification products.

### 2.14 Assessment of cell apoptosis

Cells were cultured in 6 cm dishes and treated with 0.75 μM DOX for 24 h. Following treatment, cells were harvested by trypsinization (EDTA-free) and subjected to apoptosis analysis using an annexin V-FITC/propidium iodide (PI) dual-staining kit (Elabscience, China) according to the manufacturer’s instructions. Quantitative assessment of apoptotic cells was performed by flow cytometry.

### 2.15 Oxidative stress evaluation

To assess oxidative stress, intracellular ROS were detected using 10 μM

DCFH-DA probe (Applygen, China), while mitochondrial superoxide levels were measured with 100 nM MitoSOX™ Red (Thermo Fisher Scientific, USA). Cells were incubated with the respective probes at 37°C for 30 min, washed three times with PBS, and imaged by using laser-scanning confocal microscopy.

Additionally, lipid peroxidation was evaluated by measuring malondialdehyde (MDA) content, and antioxidant capacity was determined via superoxide dismutase (SOD) activity (Beyotime, China). After cell lysis and protein quantification (BCA assay), reaction products were analyzed spectrophotometrically using a microplate reader.

### 2.16 Mitochondria isolation

Mitochondrial and cytosolic fractions were isolated using a commercial mitochondria isolation kit (Beyotime, China) according to the manufacturer’s instructions. Briefly, harvested cells were centrifuged (600 × g, 6 min, 4°C), resuspended in ice-cold isolation buffer, and incubated on ice for 15 min. Following homogenization, the lysate was centrifuged (600 × g, 10 min, 4°C) to remove nuclei and cellular debris. The supernatant was then subjected to high-speed centrifugation (12,000 × g, 10 min, 4°C) to obtain the mitochondrial pellet, while the resulting supernatant was collected as the cytosolic fraction.

### 2.17 Mitophagy assessment via mitochondria-autophagosome colocalization

To evaluate mitophagy dynamics, H9C2 cardiomyocytes were transiently transfected with RFP-LC3B plasmid (GeneChem, China) using Lipofectamine™ 3000 reagent for 24 h. Transfected cells were then treated with 0.75 μM DOX for 24 h to induce mitochondrial stress. Mitochondria were labeled with 100 nM MitoTracker Green (Invitrogen, USA) for 30 min at 37°C, followed by PBS washes. Fluorescent images were captured using laser-scanning confocal microscopy.

### 2.18 Molecular docking analysis

The three-dimensional structure of NRF1 was obtained from the Protein Data Bank (PDB ID: 8K4L), while the CARP structure was predicted using AlphaFold2 with default parameters. Both structures were prepared using AutoDockTools-1.5.7 through the following steps: (1) removal of crystallographic water molecules, (2) addition of polar hydrogen atoms, and (3) optimization of protonation states. Protein-protein docking simulations were performed using HDock (http://hdock.phys.hust.edu.cn). The resulting complexes were visualized and analyzed using PyMOL (https://pymol.org), with CARP and NRF1 represented as deep blue and cyan cartoon models, respectively. Key binding site residues were highlighted using stick models in corresponding colors to illustrate potential interaction interfaces.

### 2.19 Immunofluorescence analysis of CARP and NRF1 Colocalization

Cells were fixed with 4% paraformaldehyde for 20 min at room temperature and permeabilized with 0.1% Triton X-100 for 10 min. After blocking with 10% goat serum for 1 h at room temperature, cells were incubated with primary antibodies targeting CARP and NRF1 (4°C, overnight). Following PBS washes, cells were incubated with fluorophore-conjugated secondary antibodies for1 h. Nuclei were counterstained with DAPI-containing antifade mounting medium, and high-resolution images were acquired using laser-scanning confocal microscopy.

### 2.20 Co-immunoprecipitation (Co-IP) Analysis

To investigate protein-protein interactions, we performed co-immunoprecipitation using the protein A/G immunomagnetic bead kit (Abbkine, China) according to the manufacturer’s instructions. Briefly, H9C2 cardiomyocytes were lysed in ice-cold buffer (containing protease inhibitors) to prepare total protein extracts. A 50 μl aliquot of each lysate was saved as input control. For each reaction, 1 mg of protein lysate was pre-cleared by incubation with 20 μl magnetic beads at 4°C for 1 h to reduce nonspecific binding. The pre-cleared lysates were then immunoprecipitated with either 2 μg anti-CARP antibody or control IgG antibody for 30 mins, followed by overnight incubation at 4°C with continuous rotation. Protein-antibody complexes were captured by adding fresh magnetic beads and incubating for 2 h at 4°C. After three stringent washes with cold lysis buffer then analyzed by Western blot to identify interacting partners.

### 2.21 Construct truncated plasmids and identify the interaction domains

Full-length and domain-deleted constructs of CARP and NRF1 were obtained from Tsingke Biotechnology (Beijing, China). Flag-tagged full-length and truncated CARP plasmids (CARP Delta 1-150aa, CARP Delta 151-319aa), as well as Myc-tagged full-length and truncated NRF1 plasmids (NRF1 Delta 1-108aa, NRF1 Delta 109-304aa, NRF1 Delta 305-534aa), were cloned into the pcDNA3.1+ vector. The constructed plasmids were then transfected into HEK-293T cells. 48 h post-transfection, protein-protein interactions were assessed by Co-IP followed by Western blot analysis to map the critical interacting domains between CARP and NRF1.

### 2.22 siRNA-mediated NRF1 gene knockdown

Gene silencing experiments were performed using commercially synthesized control siRNA and NRF1-specific siRNA (Beijing, China). H9C2 cardiomyocytes were seeded in 6-well plates and transfected using lipofectamine™ 3000 reagent (Thermo Fisher Scientific, USA) in Opti-MEM medium. Transfection efficiency was evaluated 48 h post-transfection by qPCR and Western blot analysis. The PINK1-targeting siRNA sequences were: 5’-GCCCAGATGTCGTCTCAAA-3’ (Sense) and 5’-UUUGAGACGACUACUGGGC -3’ (Antisense). The NRF1-targeting siRNA sequences were: 5’-GACACGGUUGCUUCGGAAA-3’ (Sense) and 5’-UUUCCGAAGCAACCGUGUC-3’ (Antisense).

### 2.23 Statistical analysis

All quantitative data are presented as mean ± SEM unless otherwise specified. Statistical analyses were performed using GraphPad Prism 9.0 (GraphPad Software). For comparison between two groups, we applied either Student’s t-test (for normally distributed data) or Mann-Whitney U test (for non-parametric data). Multiple group comparisons were conducted using one-way ANOVA followed by appropriate post-hoc tests. Categorical variables were analyzed using Chi-square. A *P* value of < 0.05 was considered statistically significant.

## 3 Results

### 3.1 DOX downregulates CARP expression in cardiac tissues

To investigate CARP’s role in doxorubicin DIC, we established a murine model of DOX-induced cardiomyopathy (Fig. 1A). Histopathological analysis revealed pronounced myocardial structural disarray and increased collagen deposition in DOX-treated mice compared with controls (Fig. 1B), confirming successful induction of cardiac injury. Strikingly, both qPCR and Western blot demonstrated that DOX administration significantly suppressed CARP expression at the mRNA and protein levels in cardiac tissues (Fig. 1C, D). These findings strongly suggested that CARP downregulation might be involved in the pathogenesis of DIC.

**Fig 1.**
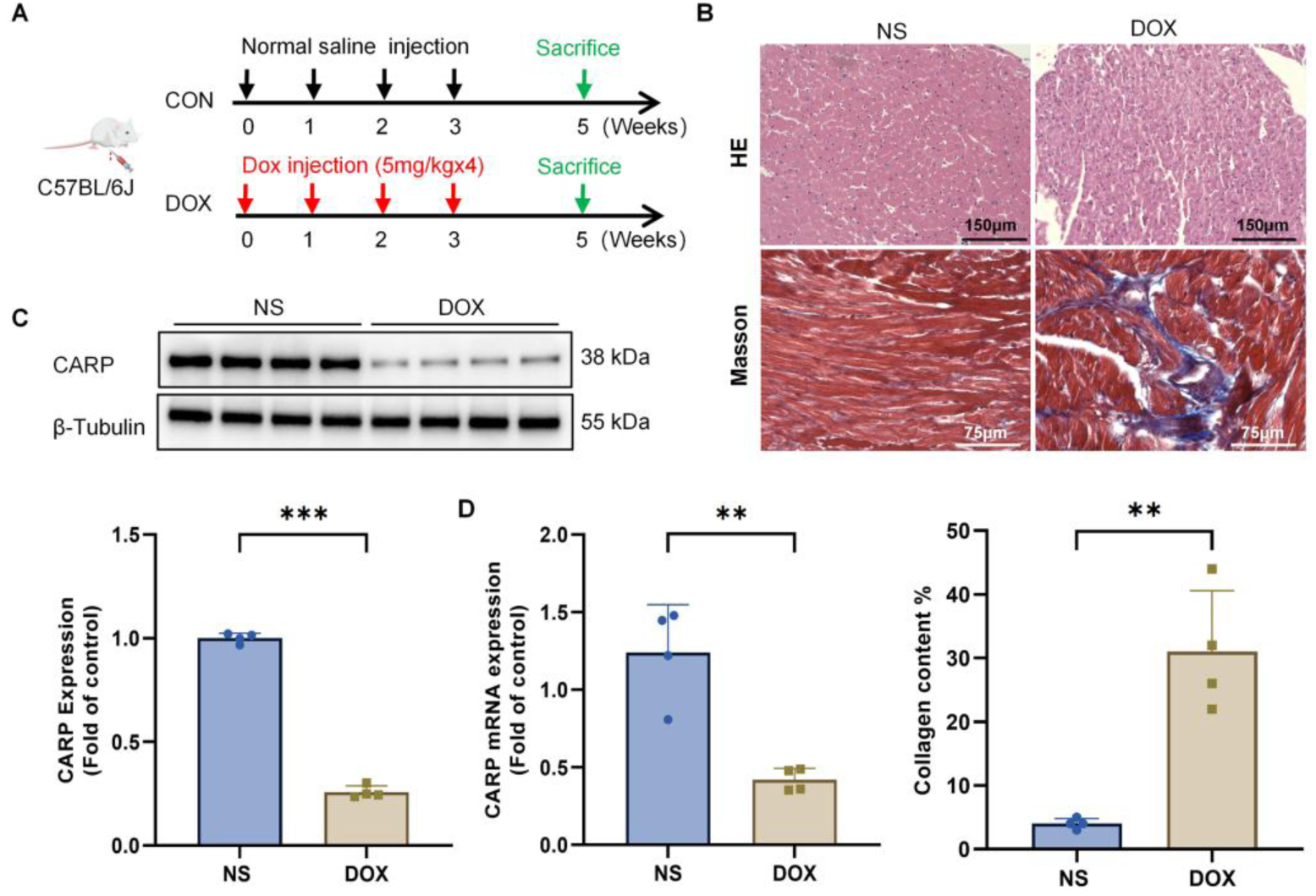
DOX induces cardiac injury and suppresses CARP expression in mice. **(A)** Experimental timeline and design of DIC in WT mice. **(B)** Representative histological analyses of myocardial tissues demonstrate: histopathological alterations in H&E-stained sections (scale bar = 150 µm) (n = 4).; collagen fiber deposition visualized by Masson’s trichrome staining (scale bar = 75 µm) (n = 4). **(C)** Western blot analysis and densitometry of CARP protein expression (n = 4). **(D)** qPCR analysis of CARP mRNA levels in cardiac tissue (n = 4). Data are mean ± SEM. Statistical significance was assessed by two-tailed unpaired t-tests: \**p* < 0.05, ** *p* < 0.01, *** *p* < 0.001; ns, not significant (*p* > 0.05).

### 3.2 Cardiac-specific CARP transgene ameliorates DOX-induced cardiac dysfunction in mice

To investigate CARP’s cardioprotective role in DIC, we generated cardiac-specific CARP-Tg mice under the α-MHC promoter (Fig. S1A). Genotypic analysis identified a 295-bp amplicon exclusive to CARP-Tg mice (Fig. S1B), qPCR and Western blot confirmed robust CARP overexpression at mRNA and protein levels in cardiomyocytes compared to WT mice (Fig. S1C, D), validating successful transgene integration.

Following DOX administration, DIC models in both WT and CARP-Tg mice were established (Fig. 2A). Echocardiographic assessment at 2 weeks post-treatment demonstrated that DOX-treated mice developed significant left ventricular (LV) systolic dysfunction, characterized by significant reductions in left ventricular ejection fraction (LVEF) and fractional shortening (LVFS), diminished left ventricular posterior wall thickness at both end-diastole (LVPWd) and end-systole (LVPWs), alongside dilated left ventricular internal dimensions during diastole (LVIDd) and systole (LVIDs). Strikingly, CARP-Tg mice exhibited preserved systolic function, with significant improvements in LVEF, LVFS, LVPWd, LVPWs, and attenuated LV dilation (Fig. 2B–H).

**Fig 2.**
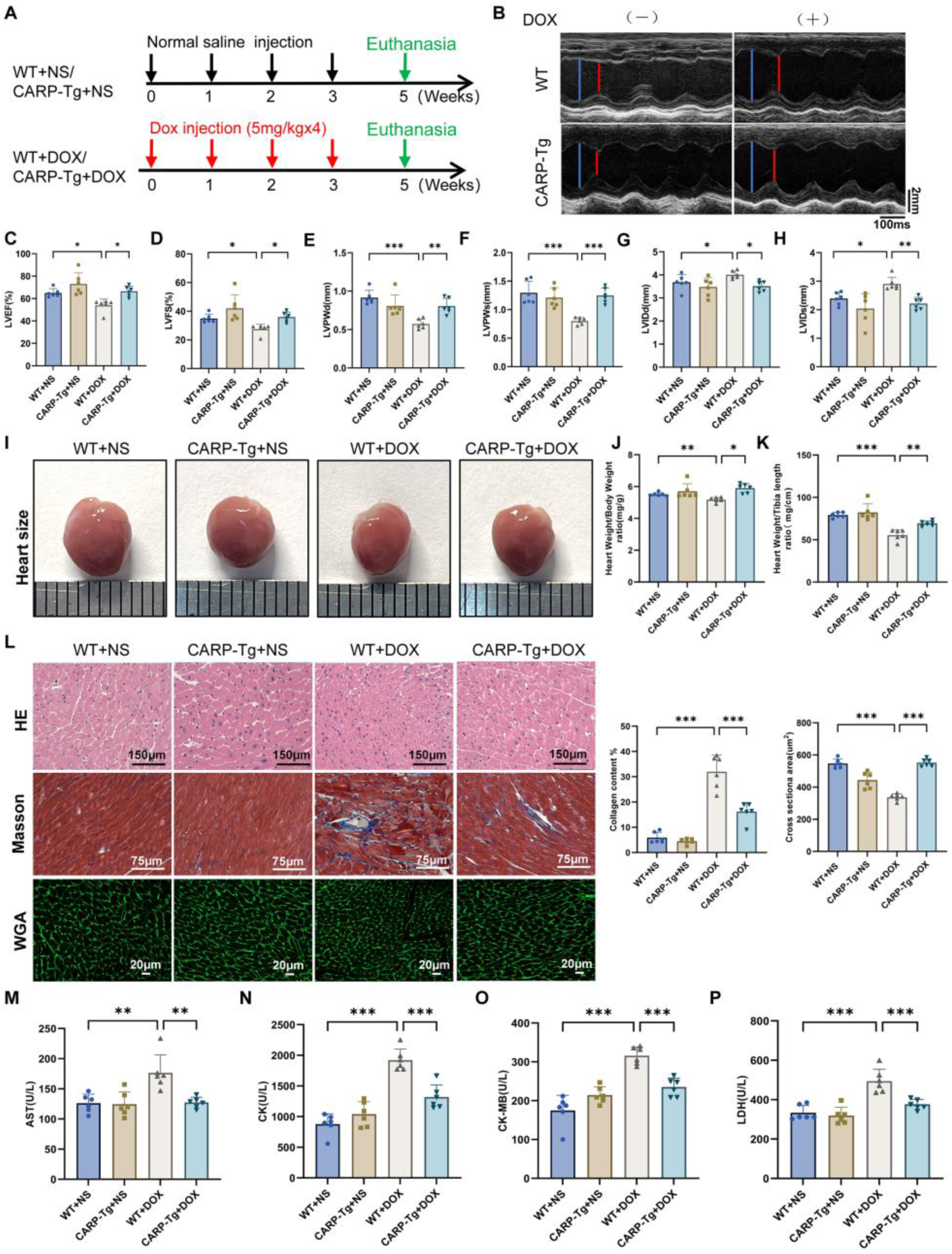
CARP attenuates DOX-induced cardiac dysfunction in mice. **(A)** Experimental schema of the DIC model. **(B)** Representative M-mode echocardiograms obtained 4 weeks post-DOX administration, with left ventricular end-systolic (LVEDs, blue) and end-diastolic (LVEDd, red) diameters indicated scale bar = 2 mm. **(C-H)** Quantitative echocardiographic assessment of cardiac function: LVEF, LVFS, LVIDs, LVIDd, LVPWd, and LVPWs (n = 6). **(I)** Macroscopic cardiac morphology across experimental groups. **(J, K)** Cardiac hypertrophy indices: HW/BW and HW/TL ratios (n = 6). **(L)** Representative histological analyses of myocardial tissues demonstrate: (top) histopathological alterations in H&E-stained sections (scale bar = 150 µm); (middle) collagen fiber deposition visualized by Masson’s trichrome staining (scale bar = 75 µm); and (bottom) cardiomyocyte cross-sectional areas identified by wheat germ agglutinin (WGA) staining (scale bar = 20 µm), with corresponding quantitative analyses shown in adjacent panels (n = 6). **(M-P)** Serum biomarkers of cardiac injury: AST, CK, CK-MB isoenzyme, and LDH levels (n = 6). Data represent mean ± SEM. Statistical significance was determined by one-way ANOVA with Tukey’s post hoc test: \**p* < 0.05, ** *p* < 0.01, *** *p* < 0.001; ns, not significant (*p* > 0.05).

DOX-treated WT mice displayed progressive body weight (BW) loss and reduced heart weight-to-body weight (HW/BW) and heart weight-to-tibia length (HW/TL) ratios, indicating cardiac atrophy. In contrast, CARP-Tg mice mitigated BW loss and maintained HW/BW and HW/TL ratios (Fig. 2I–K; Fig. S2A, B), suggesting protection against DOX-induced wasting.

Histopathology revealed that CARP-Tg mice resisted DOX-induced structural damage, showing preserved cardiomyocyte alignment, reduced fibrosis, and larger cross-sectional areas (Fig. 2L). Consistently, collagen III protein levels accumulation in DOX-treated hearts was attenuated in CARP-Tg mice (Fig. S2C).

Serum biomarkers of myocardial injury CK, its myocardial-specific isoenzyme CK-MB, LDH, and AST were elevated in DOX-treated WT mice but significantly reduced in CARP-Tg mice (Fig. 2M–P). Together, these data demonstrate that CARP overexpression preserves cardiac function, mitigates fibrotic remodeling, and attenuates biomarker release in DIC.

### 3.3 Restoration of cardiac CARP mitigates DOX-induced mitochondrial dysfunction in murine myocardium

To elucidate CARP’s role in regulating mitochondrial dysfunction during DIC, mitochondrial morphology was assessed using TEM. DOX-treated cardiac tissues exhibited severe mitochondrial damage, including cristae fragmentation, vacuolization, sarcoplasmic reticulum dilation, and myofibril disorganization (Fig. 3A). Remarkably, CARP overexpression substantially preserved mitochondrial architecture, maintaining membrane integrity, cristae density, and reducing vacuolar degeneration (Fig. 3A). Additionally, DOX treatment impaired mitochondrial biogenesis in cardiac tissues, as shown by decreased mtDNA copy number and reduced expression of key regulators PGC-1α and NRF1 (Fig. 3B, C). CARP-Tg mice, however, effectively prevented mtDNA depletion and upregulated the expression levels of PGC-1α and NRF1. (Fig. 3B, C).

**Fig 3.**
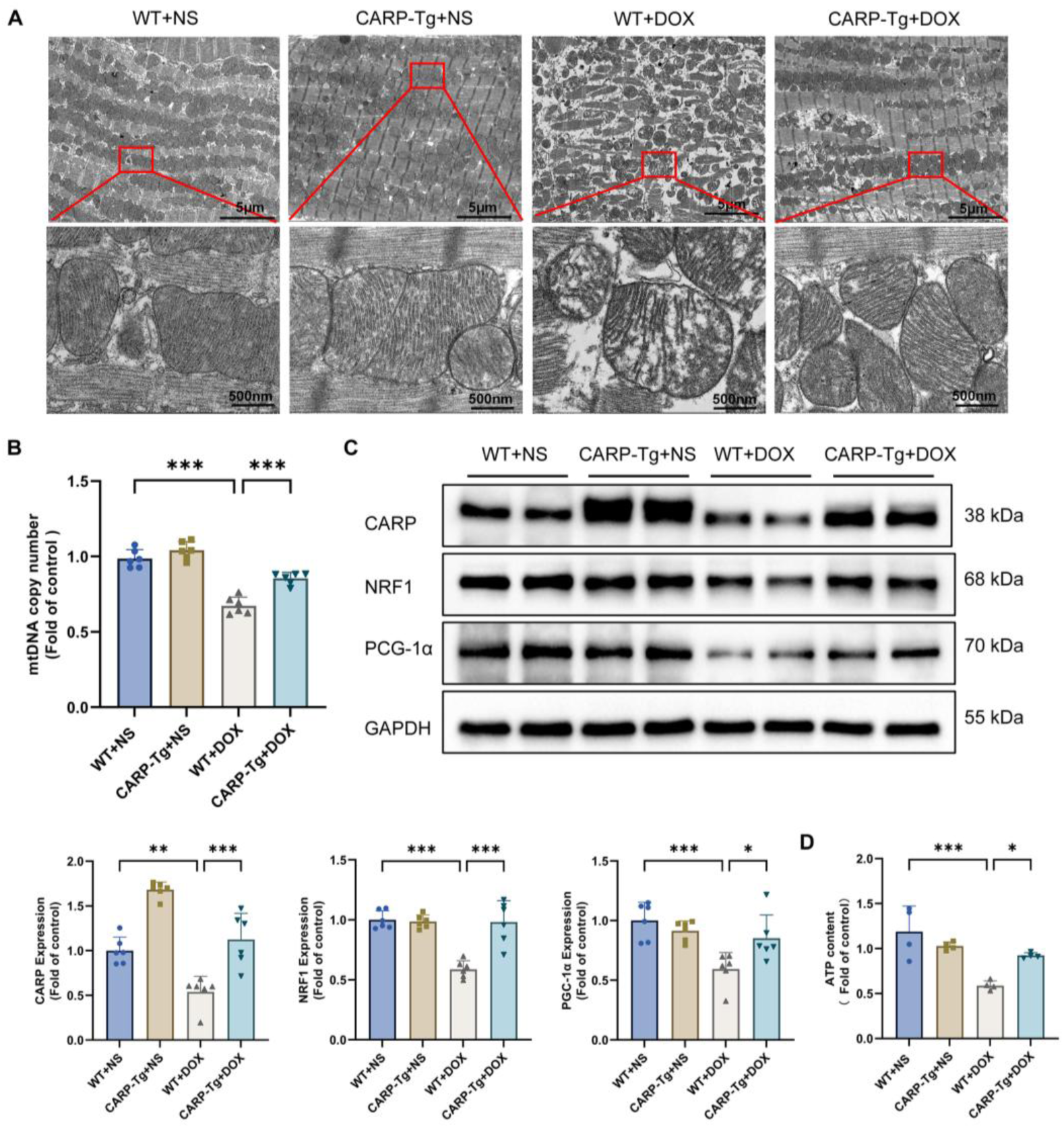
CARP ameliorates DOX-induced mitochondrial damage in mouse myocardial tissue. **(A)** TEM revealing mitochondrial ultrastructure and sarcomeric organization in myocardial tissue (scale bars provided in images). **(B)** Quantitative analysis of mtDNA copy number in cardiac tissue (n = 6). **(C)** Western blot analysis demonstrating cardiac expression levels of CARP, NRF1, and PGC-1α, with corresponding densitometric quantification (n = 6). **(D)** Myocardial ATP content measurement (n = 4). Data represent mean ± SEM Statistical significance was determined by one-way ANOVA with Tukey’s multiple comparisons test: \**p* < 0.05, ** *p* < 0.01, *** *p* < 0.001; ns, not significant (*p* > 0.05).

Additionally, DOX treatment significantly reduced myocardial ATP levels, whereas CARP-Tg mice receiving DOX demonstrated a marked increase in ATP content (Fig. 3D). Our assessment of oxidative stress markers in murine tissues revealed that DOX treatment significantly elevated MDA levels and reduced SOD activity compared with control group. Notably, CARP-Tg mice receiving DOX demonstrated significant amelioration of these oxidative stress parameters, showing decreased MDA content and restored SOD activity (Fig. S3A, B). Furthermore, TUNEL assays demonstrated CARP’s ability to reduce DOX-induced cardiomyocyte apoptosis (Fig. S3C). Collectively, these results demonstrate that CARP overexpression effectively attenuates DIC progression by ameliorating mitochondrial damage.

To further elucidate CARP’s role in mitochondrial quality control, we examined its ability to restore the balance of mitochondrial dynamics proteins disrupted by DOX treatment. Notably, CARP effectively normalized the expression levels of both DRP1 and mitochondrial fusion proteins (Mfn1/2) to physiological levels (Fig. 4A). Next, we employed TEM to examine the formation of mitophagosomes in mouse myocardial tissue. While DOX-treated hearts exhibited abundant autophagic vacuoles containing damaged mitochondria, CARP overexpression significantly reduced mitophagosome formation (Fig. 4B). Western blot analysis revealed that DOX treatment triggered excessive mitophagy, characterized by increased microtubule-associated protein light chain 3 II (LC3 II) conversion, elevated PTEN-induced putative kinase protein 1 (PINK1) and Parkin expression, and decreased p62 levels. In contrast, CARP-Tg+DOX group demonstrated reduced levels of LC3 II, PINK1, and Parkin proteins, along with elevated p62 expression (Fig. 4C). Together, these results demonstrate that CARP provides comprehensive mitochondrial protection through three synergistic mechanisms: (1) restoration of normal fission-fusion dynamics, (2) suppression of pathological mitophagy, and (3) preservation of ultrastructural integrity.

**Fig 4.**
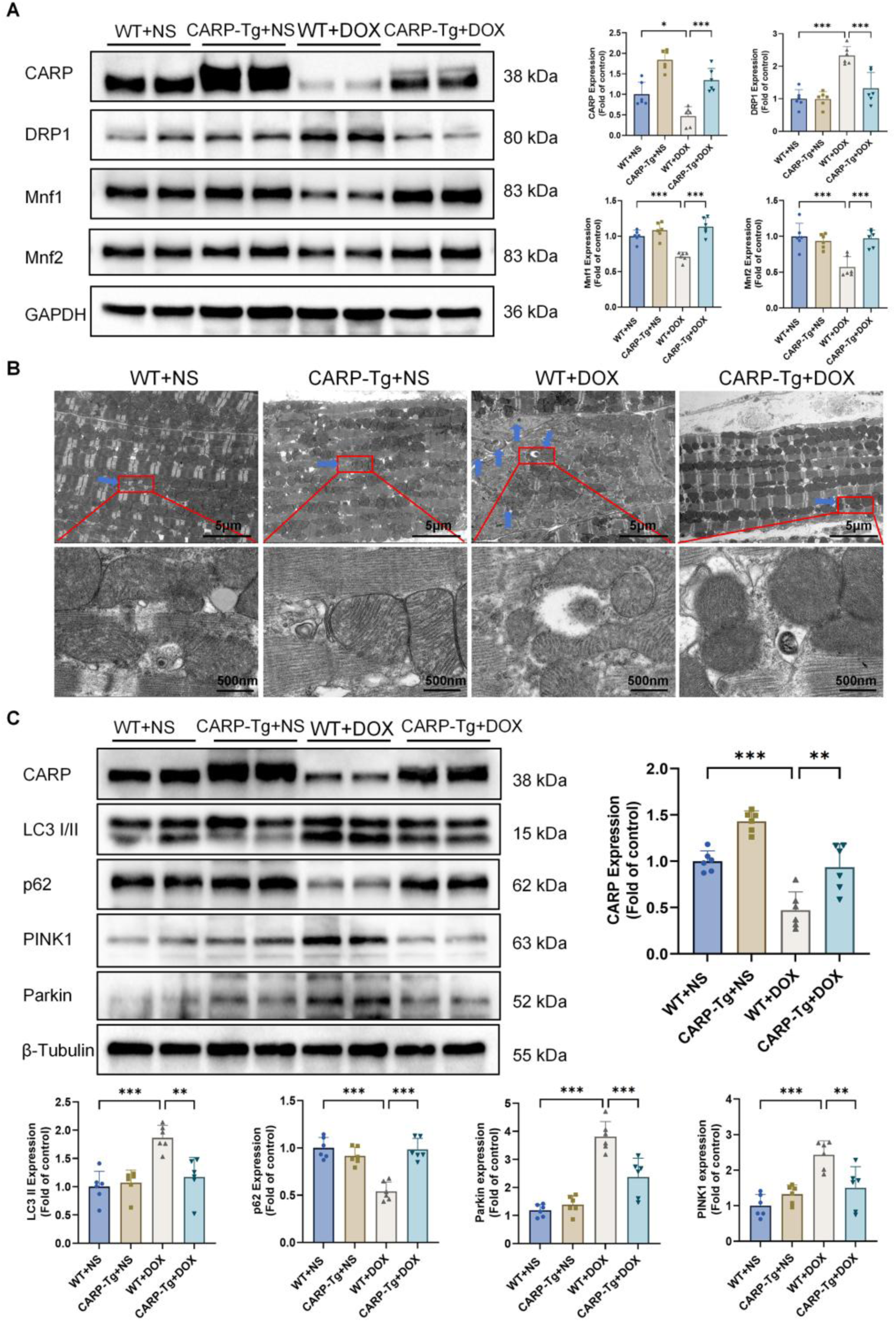
CARP overexpression modulates mitochondrial dynamics and mitophagy in DOX-treated hearts. **(A)** Western blot analysis of proteins: CARP, DRP1, Mfn1 and Mnf2 , with corresponding densitometric quantification. (n = 6). **(B)** TEM images demonstrating mitophagosome formation (blue arrows indicate autophagosomes; scale bars provided in images. **(C)** Western blot analysis of proteins: CARP, LC3 II, p62, PINK1, and Parkin, with corresponding densitometric quantification. (n = 6). Data represent mean ± SEM. Statistical significance was determined by one-way ANOVA with Tukey’s post hoc test: \**p* < 0.05, ** *p* < 0.01, *** *p* < 0.001; ns, not significant (*p* > 0.05).

### 3.4 DOX suppresses CARP expression and disrupts mitochondrial homeostasis in cardiomyocytes

To define the functional significance of CARP in DIC pathogenesis, we established a well-characterized *in vitro* model using H9C2 cardiomyocytes. Dose-response experiments revealed that DOX treatment significantly suppressed CARP expression in a concentration-dependent manner, with both mRNA and protein levels being markedly reduced (Fig. S4A, B). Next, we measured cell viability through the CCK-8 assay and noted that DOX reduced cardiomyocyte viability in a concentration-dependent manner. Notably, DOX at the concentration of 0.75 μM evoked moderate cardiomyocyte injury, with cell viability reduced by about 50% (Fig. S4C).These findings suggest that CARP suppression may represent an early molecular event contributing to DOX-induced cardiomyocyte injury, supporting its potential cardioprotective function in DIC pathogenesis.

To systematically evaluate DOX-induced mitochondrial dysfunction, we performed comprehensive analyses in H9C2 cardiomyocytes. Western blot analysis revealed that DOX treatment profoundly increased DRP1 while decreased Mfn1, Mfn2, PGC-1α and NRF1 levels (Fig. S5A, B). Using the RFP-LC3B reporter system in combination with mitochondrial-specific probes for co-localization analysis, we observed a significant increase in mitophagosome formation (yellow puncta) and mitophagosome-to-mitochondria ratio following DOX or CCCP treatment (Fig. S6A), indicating a significant increase in mitophagosome formation induced by DOX. Western blot analysis further confirmed that DOX treatment significantly increased LC3 II, PINK1, and Parkin protein levels while decreasing P62 expression (Fig. S6B), suggesting that DOX can damage cardiomyocytes and activate mitophagy.

To further elucidate the roles of mitophagy in DIC pathogenesis, we performed systematic validation through both pharmacological and genetic approaches. Initial experiments using Mdivi-1 demonstrated significantly improved cardiomyocyte viability in the Mdivi-1+DOX group compared to DOX treatment alone as assessed by CCK-8 assays (Fig. S6C). Subsequent siRNA-mediated knockdown of PINK1, a critical mitophagy regulator, was optimized through qPCR analysis, with the second siRNA construct showing about 75% knockdown efficiency and being selected for further studies (Fig. S6D). Notably, PINK1 knockdown similarly ameliorated DOX-induced cardiomyocyte viability loss (Fig. S6E). Together, these findings demonstrate that inhibiting mitophagy effectively attenuates DOX-mediated cardiomyocyte death.

To verify the effect of DOX on mitophagy flux, we employed chloroquine (CQ) co-treatment to block the autolysosomal degradation pathway. Autophagic flux analysis revealed that CQ co-treatment with DOX significantly augmented mitophagosome accumulation (Fig. S6F) and intensified LC3-II/p62 protein stabilization compared to DOX monotherapy (Fig. S6G), confirming DOX-driven autophagic flux progression rather than impairing lysosomal degradation. These findings collectively demonstrate that DOX disrupts cardiomyocyte mitochondrial homeostasis through three distinct mechanisms: (1) imbalance of fission-fusion dynamics, (2) activation of mitophagy flux, and (3) suppression of mitochondrial biogenesis.

### 3.5 CARP overexpression in cardiomyocytes confers protection through mitochondrial preservation in DIC

To elucidate the cardioprotective mechanisms of CARP in DIC pathogenesis, we generated stable CARP-overexpressing H9C2 cardiomyocytes through lentiviral transduction of Flag-tagged CARP. Quantitative analysis confirmed successful CARP overexpression at both transcriptional and translational levels compared to vector controls (Fig. S7A, B). Western blot with Flag antibody further validated expression of the CARP-Flag fusion protein (Fig. S7C). Functional characterization demonstrated that CARP overexpression significantly mitigated DOX-induced cytotoxicity, as evidenced by restored cell viability (Fig. S8A). Furthermore, CARP exhibited robust antioxidant activity, reducing both cytosolic and mitochondrial ROS accumulation (Fig. 5A), decreasing lipid peroxidation (MDA content, Fig. S8B), and enhancing SOD activity (Fig. S8C). We next examined CARP’s effects on mitochondrial functions. Relative to DOX-treated controls, CARP overexpression promoted the upregulation of mitochondrial biogenesis parameters, demonstrating increase in mtDNA content, and PGC-1α and NRF1 elevation in protein expression (Fig. 5B, C). CARP preserved mitochondrial energetics by maintaining ATP production (Fig. 5D) and MMP (Fig. 5E) in DOX-challenged cells. Flow cytometric analysis revealed CARP’s potent anti-apoptotic effects, significantly attenuating DOX-induced apoptosis rate (Fig. 5F). Western blot analysis further confirmed these findings, showing CARP overexpression reduced both the cleaved caspase-3 to caspase-3 ratio and Bax to Bcl-2 ratio (Fig. 5G).

**Fig 5.**
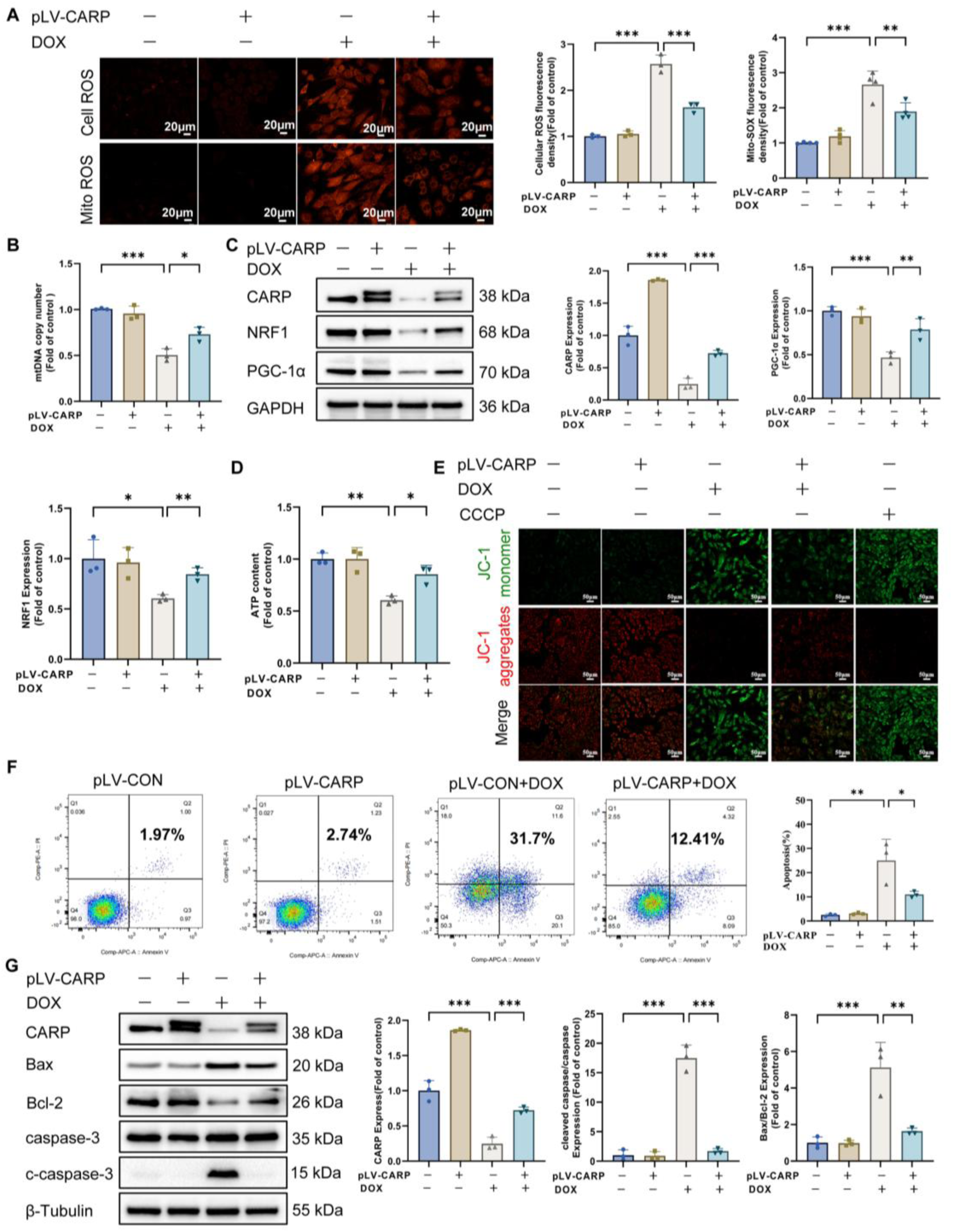
CARP overexpression preserves mitochondrial function in DOX-treated cardiomyocytes. **(A)** Representative fluorescence images and quantitative analysis of intracellular and mitochondrial ROS levels using DHE (dihydroethidium) and MitoSOX probes, scale bar = 20 μm (n = 3). **(B)** mtDNA copy number analysis (n = 3). **(C)** Western blot analysis of mitochondrial dynamics regulators (DRP1, Mfn1, Mnf2) and biogenesis factors (PGC-1α, NRF1) with corresponding densitometry (n = 3). **(D)** Cellular ATP content measurement (n = 3). **(E)** MMP assessment by JC-1 staining, scale bar=50 μm (n = 3). **(F)** quantification of apoptotic cells by flow cytometry (n = 3). **(G)** Western blot analysis of apoptosis markers (Bax, Bcl-2, c-caspase-3, and caspase-3) with corresponding densitometry (n = 3). Data represent mean ± SEM. Statistical significance was determined by one-way ANOVA with Tukey’s post hoc test: \**p* < 0.05, ** *p* < 0.01, *** *p* < 0.001; ns, not significant (*p* > 0.05).

To comprehensively characterize CARP’s regulatory role in mitochondrial quality control, we systematically examined its effects on mitochondrial dynamics and mitophagy. Western blot demonstrated that CARP overexpression effectively counteracted DOX-induced perturbations in mitochondrial dynamics proteins, decreasing DRP1 protein levels while restoring physiological levels of fusion mediators Mfn1 and Mfn2 (Fig. 6A). Using quantitative confocal microscopy with RFP-LC3b and MitoTracker co-staining, we observed that CARP overexpression significantly reduced both the absolute number of mitophagosomes (yellow puncta) and their relative proportion to total mitochondria in DOX-treated cardiomyocytes (Fig. 6B). Western blot confirmed these findings, showing CARP-mediated suppression of DOX-induced LC3-II conversion, PINK1/Parkin accumulation, and p62 degradation (Fig. 6C). Subcellular fractionation studies revealed that CARP overexpression reduced mitochondrial-associated LC3-II, PINK1, and Parkin while increasing mitochondrial p62 levels (Fig. 6D). Further supporting its role in modulating autophagic flux, CARP attenuated chloroquine-induced LC3 II accumulation and p62 depletion (Fig. S9A). Together, these actions preserve mitochondrial network integrity and function in the face of DOX-induced stress.

**Fig 6.**
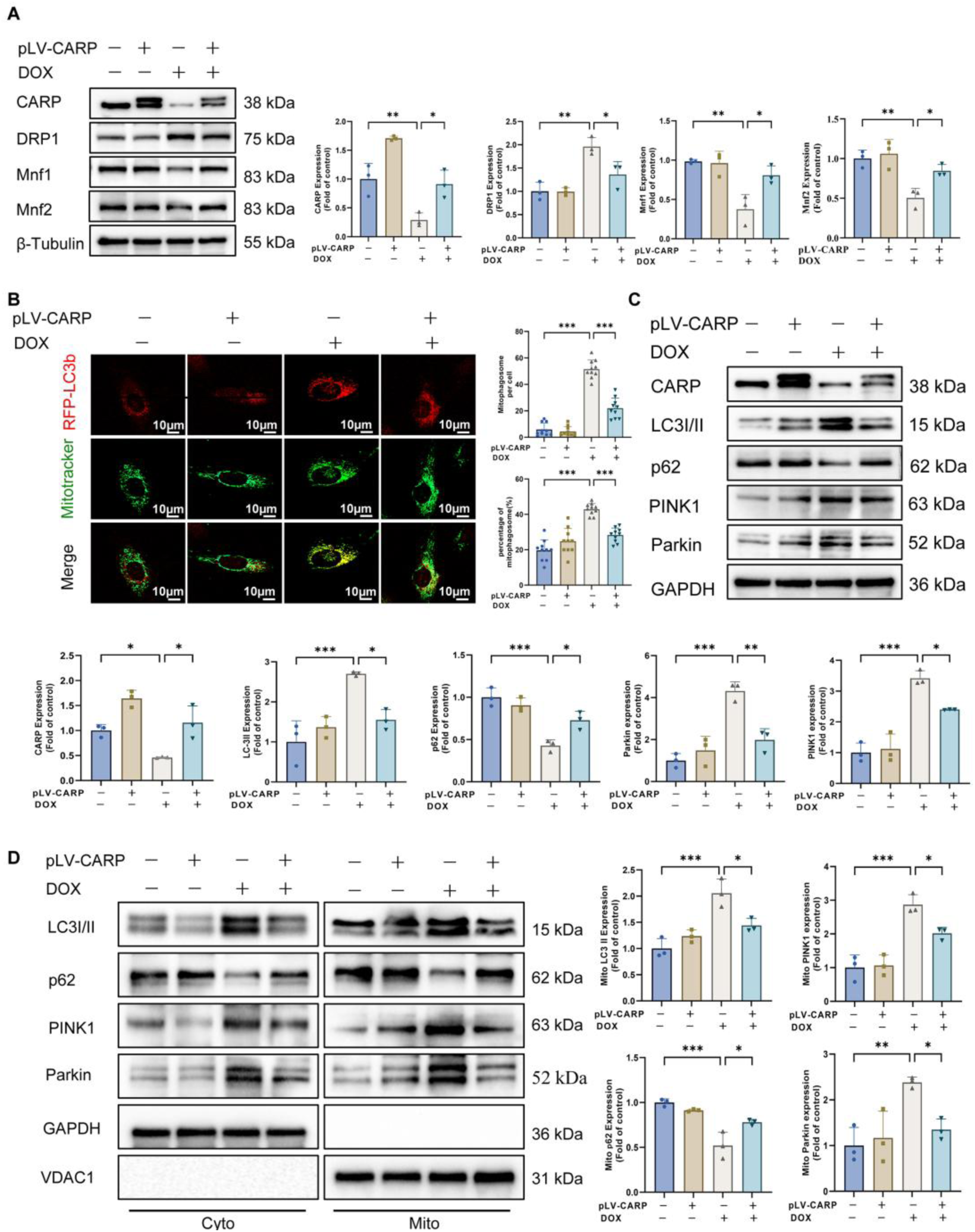
CARP overexpression attenuates DOX-induced mitochondrial fission and autophagy while enhancing fusion in cardiomyocytes. **(A)** Western blot analysis of key mitochondrial dynamics regulators (DRP1, Mfn1, Mfn2) in H9C2 cells, with representative images and quantitation (n = 3). **(B)** Fluorescent reporter system (RFP-LC3b) labeling autophagosomes. After treatment with 0.75 μM DOX for 24 hour, MitoTracker Green was used to label mitochondria. Representative confocal microscopy images and quantitative analysis of mitochondrial-autophagosome colocalization are shown, scale bar = 10 μm (n = 10 cells/group). **(C)** Western blot analysis of mitophagy proteins (LC3-II, P62 PINK1, Parkin) with representative images and quantitation (n = 3). **(D)** Representative Western blot images and quantitative analysis of LC3-II, P62, PINK1, and Parkin proteins in the cytoplasmic and mitochondrial fractions of cells (n = 3). Data represent mean ± SEM. Significance was determined by one-way ANOVA with Tukey’s post-hoc test: \**p* < 0.05, ** *p* < 0.01, *** *p* < 0.001; ns, not significant (*p* > 0.05).

### 3.6 CARP interacts with NRF1 to mediate mitochondrial protection

Next, we define the mechanism by which CARP confers cardioprotection in DIC. Prior mass spectrometry-based studies indicate that CARP may physically interacts with NRF1 ^[36]^. We hypothesize that CARP regulates mitochondrial homeostasis and thereby exerts cardioprotective effects via this interaction. Molecular docking simulations using HDock revealed a high-affinity interaction between CARP and NRF1, with PyMOL structural analysis demonstrating that their binding interface predominantly involves the N-terminal domains of both proteins (Fig. 7A). Results of immunostaining indicated that CARP and NRF1 co-localized in the nucleus (Fig. 7B). The physical interaction was confirmed by Co-IP assays in H9C2 cells, where endogenous CARP and NRF1 reciprocally interacted (Fig. 7C, D). To identify the interaction domain of these two proteins, we generated a series of truncated CARP and NRF1 constructs (Fig. 7E) and performed exogenous Co-IP in HEK293T cells. These experiments definitively identified the N-terminal regions (amino acids 1–150 of CARP and 1–108 of NRF1) as critical for their binding (Fig. 7F). These findings establish CARP as an interactor of NRF1, providing a mechanistic basis for its role in regulating mitochondrial function during DIC.

**Fig 7.**
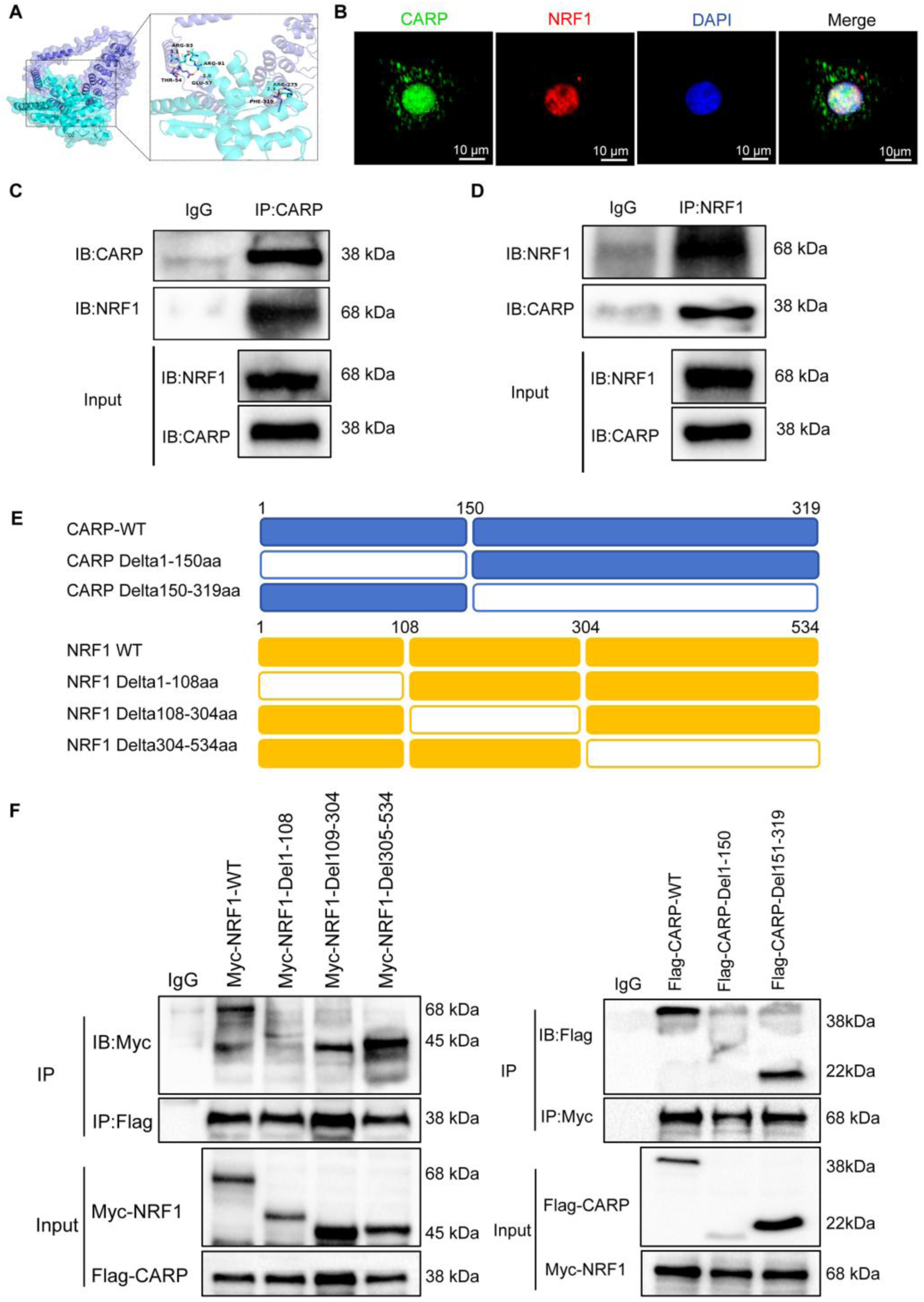
The interaction between CARP and NRF1 proteins. **(A)** Structural modeling reveals the predicted interaction interface between CARP (blue) and NRF1 (cyan), with key contact domains highlighted in the cartoon representations. **(B)** Representative immunofluorescence images showing co-localization of CARP (green) and NRF1 (red) in H9C2 H9C2 cells, scale bar = 10 μm. **(C-D)** Co-IP assays demonstrating endogenous interaction between CARP and NRF1 in H9C2 cells. **(E)** Schematic diagram of domain deletion of CARP and NRF1 for the following experiments. **(F)** Domain mapping analysis: HEK293T cells were co-transfected with Flag-tagged CARP truncations and Myc-tagged NRF1 truncations for 48 h, followed by Co-IP to identify interacting domains.

### 3.7 CARP attenuates DOX-induced mitochondrial dysfunction via NRF1 activation

To determine whether CARP exerts its mitochondrial protective effects via NRF1 activation in DIC pathogenesis, we performed NRF1 knockdown experiments in H9C2 cardiomyocytes using targeted siRNA. Among three tested siRNA constructs, siNRF1-2 demonstrated optimal knockdown efficiency, reducing mRNA expression by 70 % (Fig. S10A) and protein levels by 63% (Fig. S10B) compared to scrambled siRNA controls (si-NC). This construct was consequently employed for subsequent mechanistic investigations.

Under oxidative stress conditions, CARP overexpression alleviated DOX-induced damage, as evidenced by reducing ROS generation in both cytosol and mitochondria (Fig. 8A-B), increasing SOD activity (Fig. S11A) and decreasing MDA levels. This alleviation did not further present improvement after inducing NRF1 deficiency. Notably, NRF1 deficiency also attenuated CARP-mediated preservation of cardiomyocyte viability (Fig. S11C). NRF1 knockdown suppressed CARP-induced restoration of mitochondrial function, including mtDNA copy number recovery (Fig. 8C), PGC-1α upregulation (Fig. 8D), and partial rescue of ATP production (Fig. 8E). Furthermore, while CARP overexpression significantly decreased DOX-induced apoptosis (as measured by apoptotic rate, cleaved caspase-3 to caspase-3 ratio, and Bax to Bcl-2 ratio, these anti-apoptotic effects were substantially diminished upon NRF1 knockdown (Fig. 8F-G). These comprehensive data establish NRF1 as the essential mediator through which CARP preserves mitochondrial function and cellular viability in DIC.

**Fig 8.**
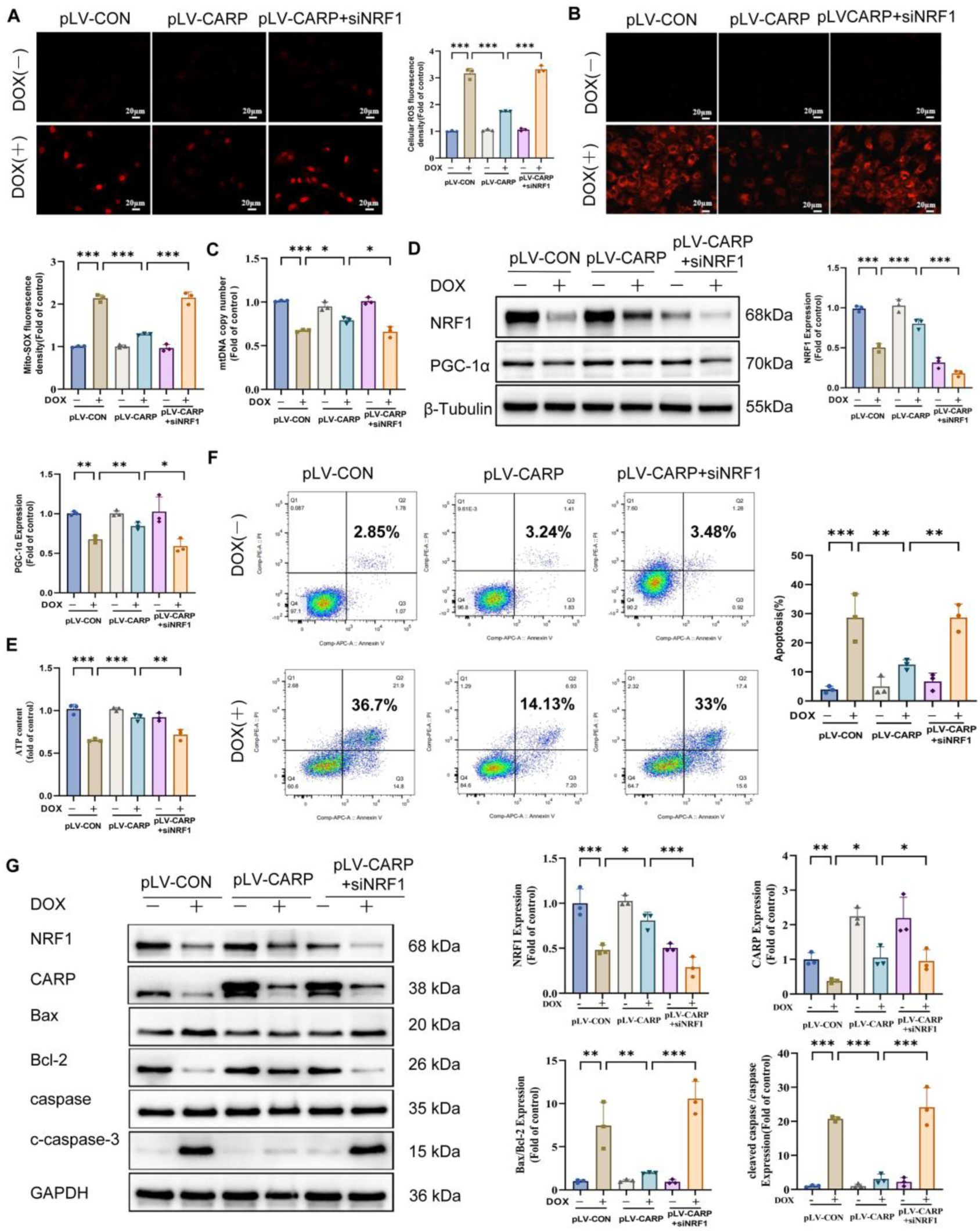
CARP mitigates DOX-induced mitochondrial dysfunction through NRF1-dependent mechanisms. (A-B) Representative fluorescence images and quantitative analysis of intracellular and mitochondrial ROS levels using DHE (dihydroethidium) and MitoSOX probes, scale bar = 20 μm (n = 3). **(C)** mtDNA copy number analysis (n = 3). **(D)** Western blot analysis of mitochondrial biogenesis factors (PGC-1α, NRF1) with corresponding densitometry (n = 3). **(E)** ATP content measurement (n = 3). **(F-G)** Western blot analysis of CARP, NRF1, Bax, Bcl-2, c-caspase-3, and caspase-3 with corresponding densitometry (n = 3). Data represent means ± SEM. Statistical significance was determined by one-way NOVA with Tukey’s post hoc test: \**p* < 0.05, ** *p* < 0.01, *** *p* < 0.001; ns, not significant (*p* > 0.05).

Our studies demonstrate that CARP orchestrates mitochondrial quality control through NRF1-mediated mechanisms. Western blot analysis demonstrated that CARP overexpression counteracted DOX-induced alterations by reducing DRP1, LC3 II, PINK1, and Parkin levels, while elevating Mfn1, Mfn2 and p62 expression. However, NRF1 knockdown significantly attenuated these CARP-mediated regulatory effects (Fig. 9A). Using RFP-LC3b and MitoTracker Green co-localization, we observed that NRF1 knockdown significantly compromised CARP-mediated suppression of DOX-induced mitophagy, evidenced by increased autophagosome formation and enhanced mitochondrial-autophagosome co-localization (Fig. 9B). Mitochondrial fractionation assays revealed that NRF1 knockdown failed to restore CARP-mediated regulation of LC3-II, p62, PINK1, and Parkin protein levels in mitochondria under DOX treatment (Fig. 9C). Furthermore, autophagic flux assays showed NRF1 knockdown impaired CARP’s suppression of DOX-induced autophagic activity (Fig. S12A). These coordinated findings establish that CARP maintains mitochondrial homeostasis through NRF1-mediated regulation of fission, fusion and mitophagy, ultimately providing cardiomyocyte protection under DOX stress.

**Fig 9.**
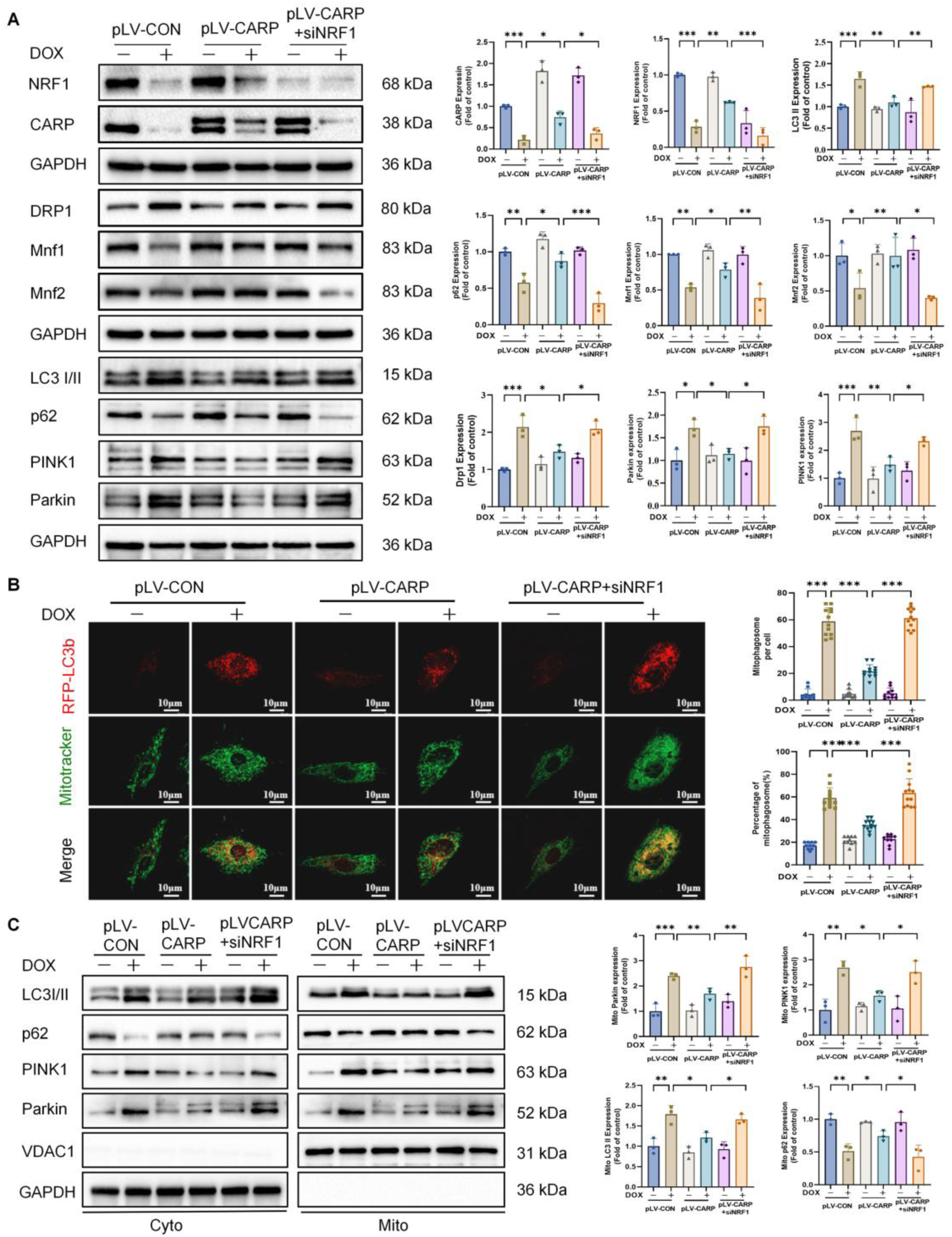
CARP attenuates DOX-induced mitochondrial dynamics dysregulation and impaired mitophagy through NRF1 signaling. **(A)** Western blot analysis of CARP, NRF1, DRP1, Mfn1, Mfn2, LC3-II, P62, PINK1and Parkin with representative images and quantitation (n = 3). **(B)** Confocal microscopy imaging and quantification of mitochondrial-autophagosome colocalization, scale bar = 10 μm (n=10 cells/group). **(C)** Representative Western blot images and quantitative analysis of LC3-II, P62, PINK1, and Parkin proteins in the cytoplasmic and mitochondrial fractions of cells (n=3). Data represent means ± SEM. Statistical significance was determined by one-way ANOVA with Tukey’s post hoc test\**p* < 0.05, ** *p* < 0.01, *** *p* < 0.001; ns, not significant (*p* > 0.05).

## 4 Discussion

Although DOX remains a cornerstone anthracycline chemotherapeutic agent for treating both pediatric and adult malignancies, its clinical utility is significantly limited by dose-dependent cardiotoxicity ^[46]^. Our study reveals a novel cardioprotective mechanism mediated by the CARP-NRF1 regulatory axis, which mitigates DIC through comprehensive maintenance of mitochondrial homeostasis (Fig. 10). We demonstrate that CARP coordinates a previously unidentified molecular defense system that preserves mitochondrial structural integrity and functional capacity during DOX challenge. These findings provide crucial insights into the molecular pathogenesis of DIC while establishing CARP as a potential therapeutic target for preventing chemotherapy-induced cardiac dysfunction.

**Fig 10.**
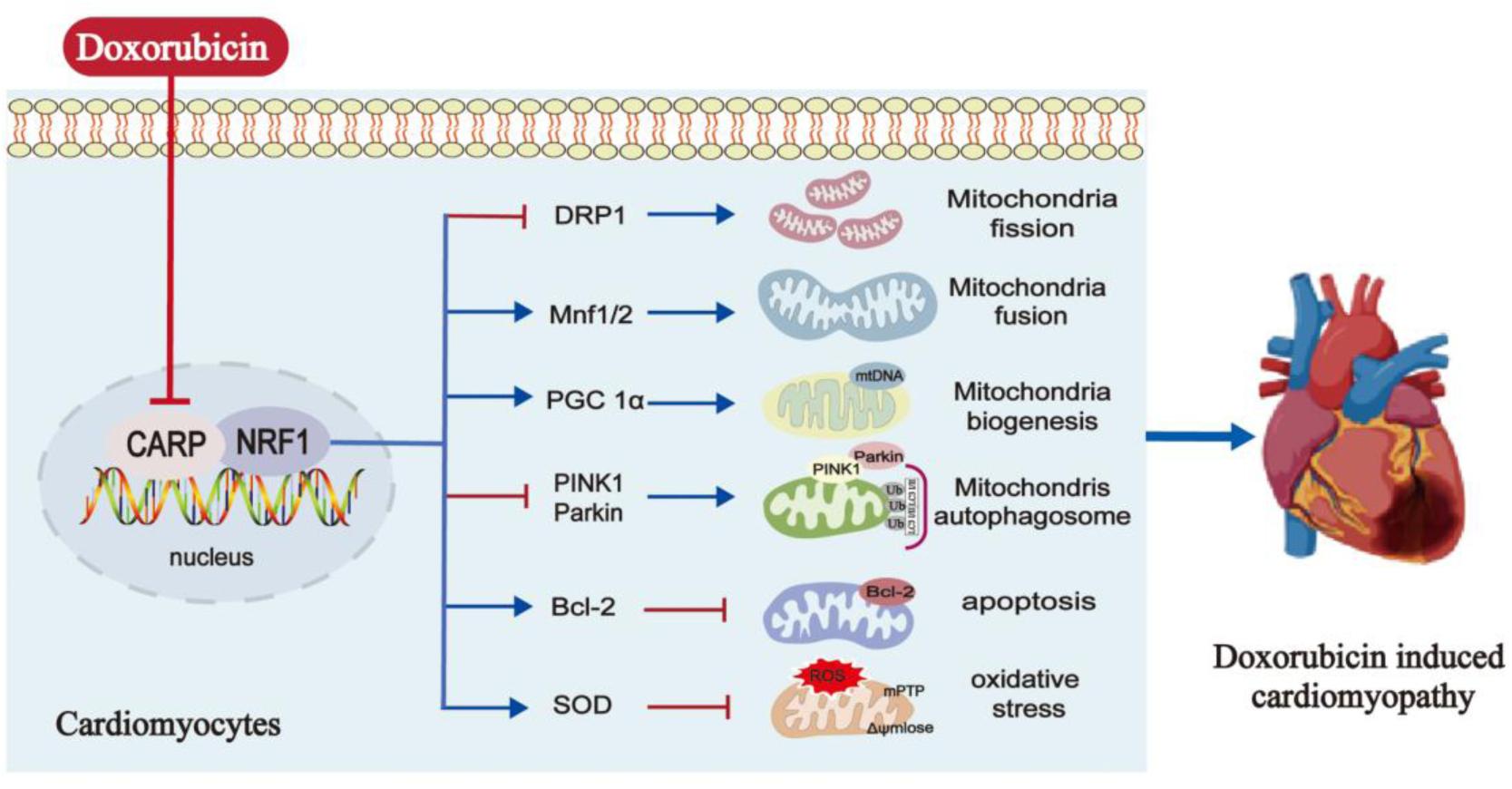
Mechanistic basis of CARP-mediated protection against DIC. This schematic illustrates how CARP overexpression protects against doxorubicin-induced cardiomyopathy (DIC) primarily through NRF1-dependent pathways. CARP exerts its cardioprotective effects by concurrently inhibiting apoptotic pathways and oxidative damage while preserving mitochondrial function via three interconnected mechanisms: (1) modulation of mitophagic flux, (2) maintenance of mitochondrial fission/fusion balance through regulation of DRP1 and mitofusins, and (3) promotion of mitochondrial biogenesis. Together, these coordinated actions effectively mitigate doxorubicin-induced cardiomyocyte damage.

### 4.1 Cardiac-specific CARP overexpression confers protection against DIC

Through the generation of CARP-Tg mice, we demonstrate that CARP overexpression effectively preserves both cardiac structure and function in DIC. Our findings reveal that CARP overexpression significantly improves left ventricular systolic performance while reducing myocardial fibrosis, underscoring its therapeutic potential against DIC. Previous studies have identified CARP as a critical regulator of sarcomere integrity, where it forms a tetrameric complex with Titin, Myopalladin, and Calpain 3/p94 to maintain cardiomyocyte structure and contractility ^[30]^. Our current work expands this understanding by demonstrating that CARP overexpression directly prevents DOX-induced myofibrillar disarray, thereby maintaining proper sarcomere organization.

The cardioprotective effects of CARP are further supported by evidence from multiple cardiac injury models. In pressure-overload-induced hypertrophy, CARP transgenic mice show significantly attenuated myocardial hypertrophy and reduced fibrosis ^[47, 48]^. Moreover, GATA4-mediated CARP upregulation during DOX treatment prevents sarcomere disorganization, whereas CARP knockdown attenuated the cardioprotective effects. ^[49]^. These consistent findings across different experimental models firmly establish CARP’s essential role in preserving contractile function and limiting cardiac injury progression.

### 4.2 CARP-mediated mitochondrial protection in DIC

Mitochondrial dysfunction represents a central pathological feature of DIC ^[50, 51]^. DOX, as a mitochondrial-targeting agent, preferentially accumulates in the mitochondrial matrix, inducing excessive ROS production ^[52, 53]^ and initiating intrinsic apoptosis through mitochondrial membrane permeabilization ^[54, 55]^. Our study reveals that CARP provides comprehensive mitochondrial protection through three synergistic mechanisms: (1) attenuation of oxidative stress via dual modulation of ROS production and endogenous antioxidant defenses (as evidenced by increased SOD activity); (2) preservation of mitochondrial bioenergetics through restoration of impaired ATP synthesis; (3) suppression of apoptosis, demonstrated by normalization of cleaved caspase-3 to caspase-3 and Bax to Bcl-2 ratios, aligning with CARP’s demonstrated ability to enhance cardiomyocyte resistance to hypoxia-induced apoptosis in myocardial ischemia models ^[35, 56, 57]^. These results position CARP as a critical regulator of cardiomyocyte survival through mitochondrial protection.

The maintenance of mitochondrial homeostasis requires precise coordination of fission-fusion dynamics and selective mitophagy. While essential for cellular physiology, dysregulation of these processes can lead to pathological mitochondrial depletion and cell death^[58, 59]^. Our data demonstrate that DOX disrupts this balance by inducing pathological hyper-fission while suppressing fusion, consistent with prior reports linking mitochondrial fragmentation to DIC progression ^[60, 61]^. Notably, CARP overexpression restores this equilibrium through dual actions: (1) inhibition of DRP1-mediated fission and (2) promotion of mitochondrial fusion.

The role of mitophagy in DIC pathogenesis remains debated^[62]^, with some studies suggesting a protective function through clearance of damaged mitochondria ^[63, 64]^, while other (including our findings) implicate excessive mitophagy in cardiac dysfunction ^[18, 51, 60, 65]^. As a central regulator of mitochondrial quality control, the PINK1/Parkin pathway has been well-documented to maintain mitochondrial homeostasis under physiological conditions ^[66]^. However, our results, consistent with previous reports ^[60, 65]^, demonstrate that DOX treatment leads to pathological activation of this pathway in cardiomyocytes, as indicated by the marked upregulation of both PINK1 and Parkin protein expression. Notably, CARP overexpression effectively suppresses this process by reducing abnormal PINK1 accumulation on mitochondria and blocking Parkin translocation to mitochondria, thereby attenuating mitophagic flux.

### 4.3 CARP orchestrates mitochondrial homeostasis via NRF1-dependent networks

Mitochondrial homeostasis necessitates precise nuclear-mitochondrial crosstalk to sustain bioenergetic competence and functional resilience ^[67, 68]^. Here, we identify CARP as a nuclear-encoded transcriptional regulator governing this essential equilibrium. Co-IP assays in H9C2 cells confirmed direct CARP-NRF1 interaction, unveiling a cardioprotective axis through coordinated transcriptional regulation.

CARP overexpression restored DOX-impaired mitochondrial bioenergetics by upregulating PGC-1α expression (a master mitochondrial biogenesis regulator ^[69]^) and enhancing mtDNA replication capacity. Notably, NRF1 knockdown compromised CARP-driven mitochondrial biogenesis, establishing NRF1 as the indispensable mediator of this pathway—consistent with its canonical role in activating PGC-1α transcription ^[70]^ and coordinating mitochondrial gene networks ^[71, 72]^. Beyond biogenesis, we delineated NRF1’s dual regulatory functions: (1) mitigating oxidative stress by amplifying CARP’s antioxidant capacity (aligned with NRF1’s redox-modulatory properties ^[41, 73]^) and (2) maintaining mitochondrial quality through fission suppression (via DRP1 transcriptional control) ^[44]^. and mitophagy optimization ^[42, 74, 75]^. Mechanistically, CARP recruits NRF1 to synchronize mitochondrial dynamics, suppressing pathological fission, mitophagy flux, and promoting fusion, while concurrently counteracting oxidative damage. The axis’s cytoprotective efficacy was further evidenced by NRF1 knockdown attenuating CARP-mediated apoptosis resistance, reinforcing NRF1’s role as the central integrator of mitochondrial-redox homeostasis ^[46, 76]^.

Notably, the cardioprotective function of NRF1 in doxorubicin-induced cardiomyopathy identified in our investigation aligns closely with emerging insights into cardiac regeneration and repair mechanisms ^[77]^. Accumulating evidence reveals that NRF1—a prominently expressed member of the Cap’n’Collar transcription factor family in cardiomyocytes—is selectively upregulated during myocardial regeneration. Genetic ablation of NRF1 disrupts transcriptional networks indispensable for cardiac repair, while its ectopic expression confers robust protection against ischemia/reperfusion injury in mature myocardium. Significantly, studies employing human induced pluripotent stem cell-derived cardiomyocyte models corroborate that NRF1 upregulation substantially attenuates DIC, reinforcing its mechanistic relevance in anthracycline-associated cardiac pathophysiology ^[77]^.

These findings position CARP-NRF1 signaling as a multilevel orchestrator of mitochondrial adaptation under cardiotoxic stress, revealing therapeutic targets to preserve myocardial function during anthracycline therapy.

## 5 Conclusion and perspectives

Our study establishes that CARP mitigates DIC by maintaining mitochondrial homeostasis through the NRF1 pathway. We present compelling evidence that CARP coordinately regulates multiple facets of mitochondrial quality control, encompassing mitophagy regulation, biogenesis enhancement, dynamics modulation, antioxidant defense, and anti-apoptotic mechanisms. These findings position CARP as a promising therapeutic target for DIC intervention.

The discovery of the CARP-NRF1 regulatory axis provides a novel therapeutic strategy against DIC. Future development of CARP-targeted agonists or small-molecule modulators of this pathway may lead to breakthrough treatments for chemotherapy-induced cardiac injury. This approach holds promise for preserving cardiac function in cancer patients undergoing anthracycline therapy.

## Author contributions

Guoda Ma and Yajun Wang designed the research. Dan Zhong, Xusan Xu, Taoshan Feng, Wensen Zhang, and Zhengqiang Luo carried out the experiments. Dan Zhong and Xusan Xu performed structure modeling. Fei Luo and Junhao Guo performed data analyses and prepared the figures. Dan Zhong and Xusan Xu wrote the original draft of the manuscript. Guoda Ma, Yajun Wang and Riling Chen revised the manuscript. The authors have read and approved the final manuscript.

## Acknowledgments

We gratefully acknowledge Professor Snezana Kojic and Professor Dragica Radojkovic (Institute of Molecular Genetics and Genetic Engineering, University of Belgrade, Serbia) for their valuable suggestions and manuscript polishing during the preparation of this work.

