## Supplementary figure and table for "CARP Protects Against Doxorubicin-induced Cardiotoxicity through NRF1-driven Mitochondrial Homeostasis": 2025-10-09Supplementary figure and table已修改.docx (from another version of this preprint: version 1, 2025-11-06)

**Table S1 Primers used for RT-PCR analysis**

| **Gene** | **Primers** | **Species** | **Sequence 5’-3’** |
| --- | --- | --- | --- |
| GAPDH | Forward | Rat | GTCCATGCCATCACTGCCACTC |
| GAPDH | Reverse | Rat | CGCCTGCTTCACCACCTTCTTG |
| NRF1 | Forward | Rat | TCGCCTGAGGACACTTCTTACG |
| NRF1 | Reverse | Rat | GCAGCAGCAGCAGCCATTC |
| ACTB | Forward | Rat | TGAGAGGGAAATCGTGCGTGAC |
| ACTB | Reverse | Rat | GGAAGAGGATGCGGCAGTGG |
| ND1 | Forward | Rat | AAGCGGCTCCTTCTCCCTACAAAT |
| ND1 | Reverse | Rat | GAAGGGAGCTCGATTTGTTTCTGC |
| ANKRD1 | Forward | Rat | AAAGTTCGAGAGGCAGAGCT |
| ANKRD1 | Reverse | Rat | GCAGTTTGTTCTCCAGTGCA |
| GAPDH | Forward | Mouse | AGTGTTTCCTCGTCCCGTAG |
| GAPDH | Reverse | Mouse | GCCGTGAGTGGAGTCATACT |
| ANKRD1 | Forward | Mouse | TGAATCCACAGCCATCCACT |
| ANKRD1 | Reverse | Mouse | CTCCTTCTCTGTCCTTGGCA |
| ACTB | Forward | Mouse | TACTGCTCTGGCTCCTAGC |
| ACTB | Reverse | Mouse | CGGACTCATCGTACTCCTGC |
| ND1 | Forward | Mouse | GGCCCATTCGCGTTATTCTT |
| ND1 | Reverse | Mouse | TCTAACGGAAGCGTGGATA |

**Table S2 Main reagents and resources used in this study**

| **Antibodies** | **Source** | **Identifier** |
| --- | --- | --- |
| Anti-CARP(G-2) | Santa Cruzsc | Catalog#365056 |
| Anti-LC3b | CellSignalingTechnology | Catalog#43566T |
| Anti-P62 | MBL | Catalog#PM045 |
| Anti-PINK1 | Proteintech | Catalog#23274 |
| Anti-Parkin | HuaBio | Catalog#ET1702 |
| Anti-DRP1 | Abcam | Catalog#ab184247 |
| Anti-Mnf1 | Proteintech | Catalog#13798 |
| Anti-Mnf2 | Abcam | Catalog#ab124773 |
| Anti-PGC 1α | Proteintech | Catalog#66369 |
| Anti-NRF1 | Abcam | Catalog#ab175932 |
| Anti-CollagenIII | Abcam | Catalog#ab184993 |
| Anti-GAPDH | Affinity | Catalog#AF7021 |
| Anti-β-Tubulin | Proteintech | Catalog#10094 |
| Anti-VDAC1 | Proteintech | Catalog#55259 |
| Anti-Myc-tag pAb | Proteintech | Catalog#60003 |
| Anti-DDDDK-tag mAb | MBL | Catalog#M185-3 |

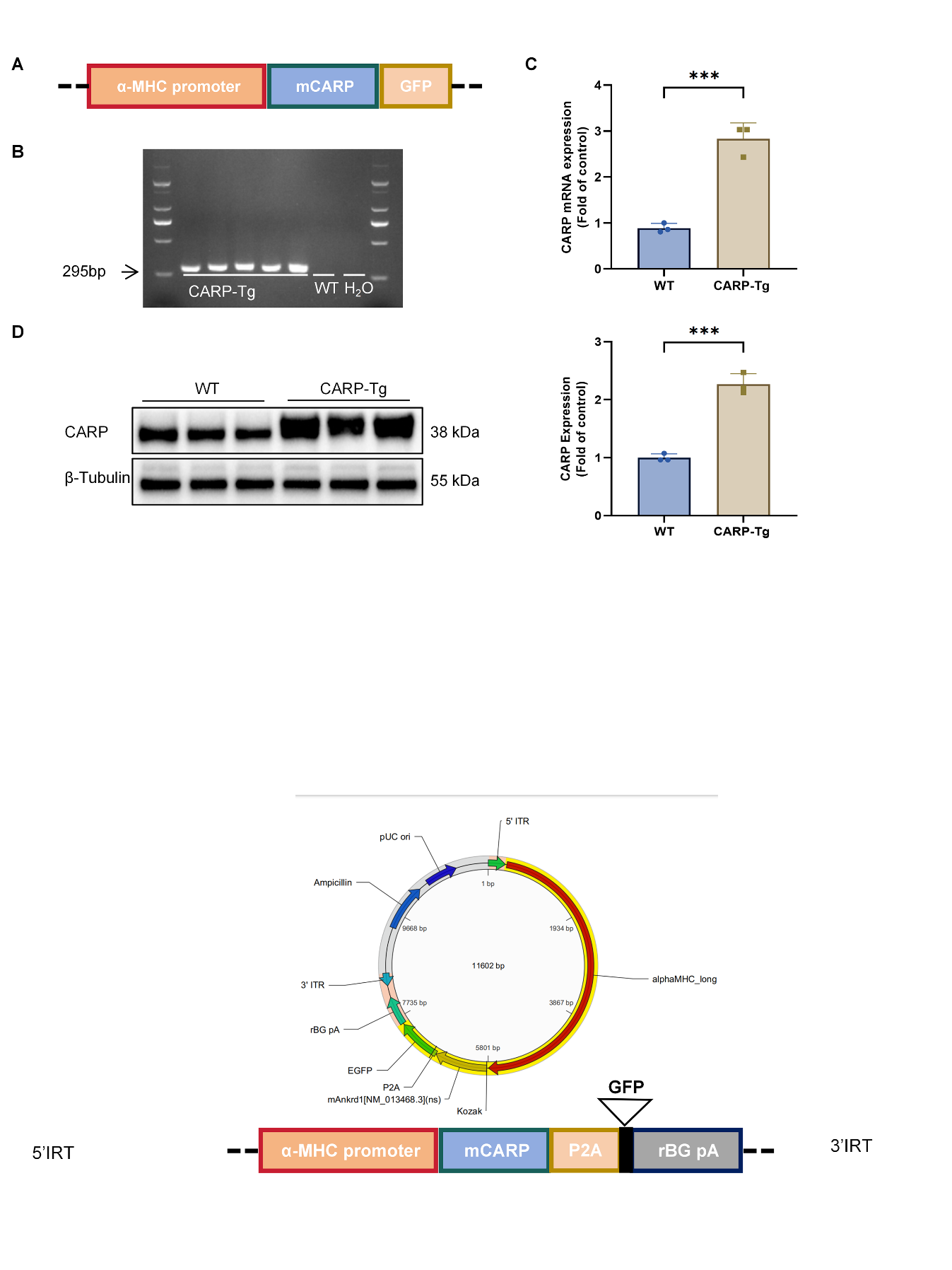

**Fig S1 Generation of cardiac-specific CARP transgenic mice.**

**(A)** Schematic diagram of the CARP transgenic construct. **(B)** Genotyping identification of CARP transgenic mice by agarose gel electrophoresis of the 295 bp PCR product. **(C)** qPCR analysis of CARP mRNA levels in cardiac tissue (n = 3). **(D)** Western blot analysis and densitometry of CARP protein expression (n = 3). Data are mean ± SEM. Statistical significance was assessed by two-tailed unpaired t-tests: **p* < 0.05, ** *p* < 0.01, *** *p* < 0.001; ns, not significant (*p* > 0.05).

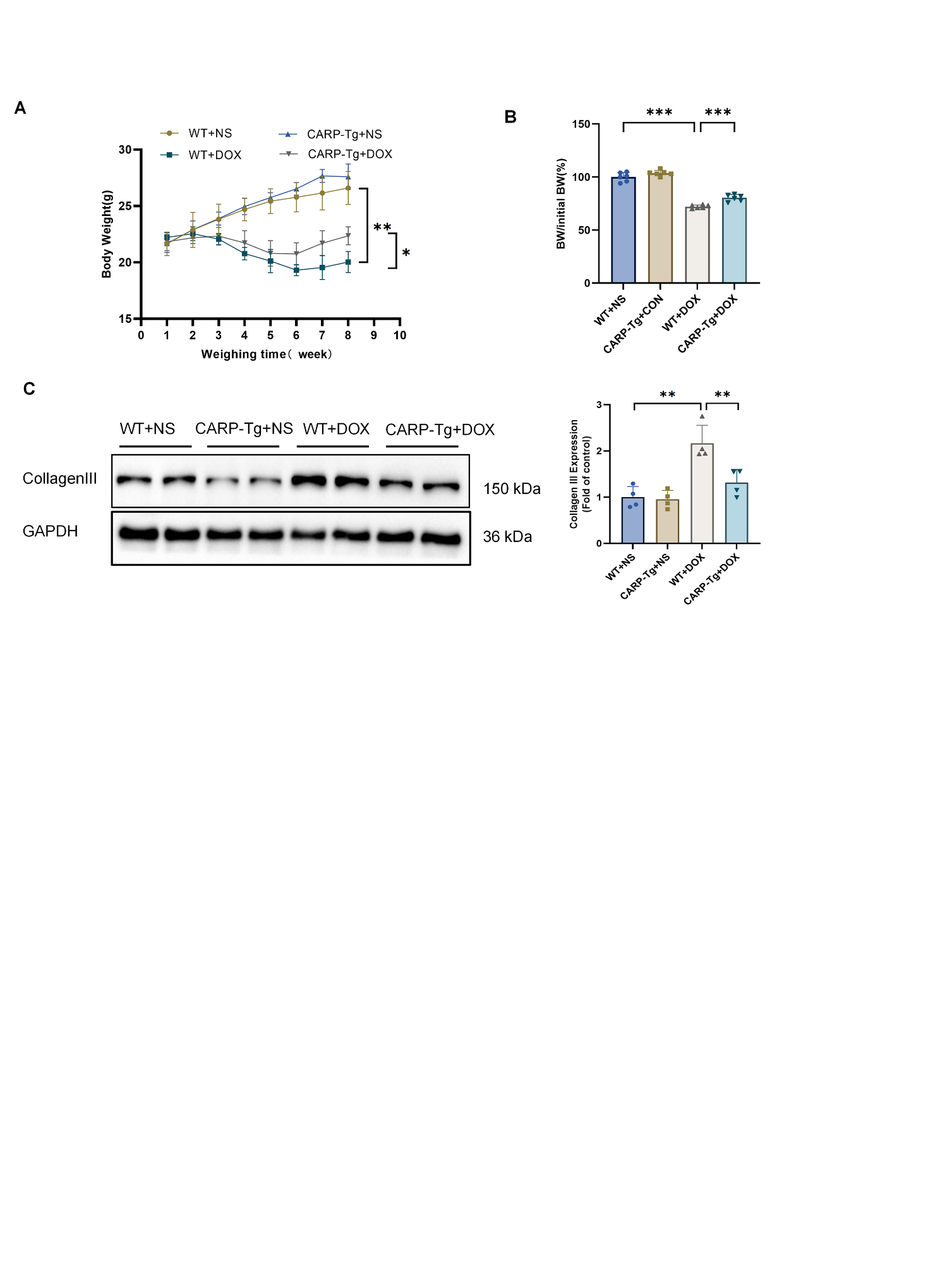

**Fig S2 CARP overexpression alleviates DOX-induced body weight loss and cardiac fibrosis in mice.**

**(A)** Weekly body weight change curves (n = 6). **(B)** BW/initial BW ratios (n = 6). **(C)** Western blot analysis demonstrating cardiac expression levels of Collagen III, with corresponding densitometric quantification (n = 4). Data represent mean ± SEM. Statistical significance was determined by one-way ANOVA with Tukey's post hoc test: **p* < 0.05, ** *p* < 0.01, *** *p* < 0.001; ns, not significant (*p* > 0.05).

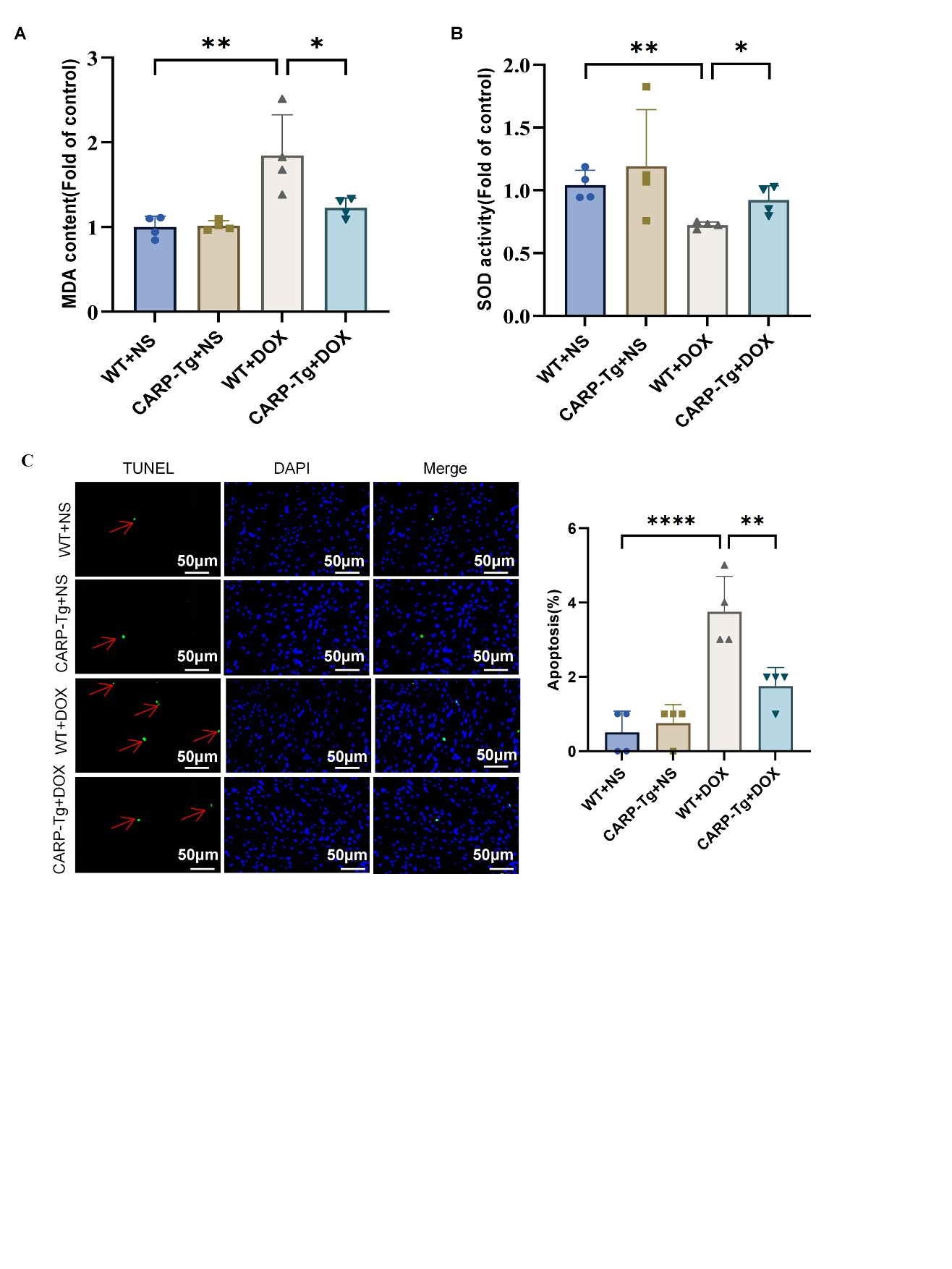

**Fig S3 CARP overexpression attenuates DOX-induced oxidative stress and apoptosis**.

**(A)** ROS levels in cardiac tissues detected by DHE probe (n = 4). **(B)** MDA content in cardiac tissues (n = 4). **(C)** SOD activity in cardiac tissues (n = 4). **(D)** Representative TUNEL staining images and quantitative analysis of apoptosis in myocardial tissues, scale bar = 50 μm (n = 4). Data represent mean ± SEM. Statistical significance was determined by one-way ANOVA with Tukey's post hoc test: **p* < 0.05, ** *p* < 0.01, *** *p* < 0.001; ns, not significant (*p* > 0.05).

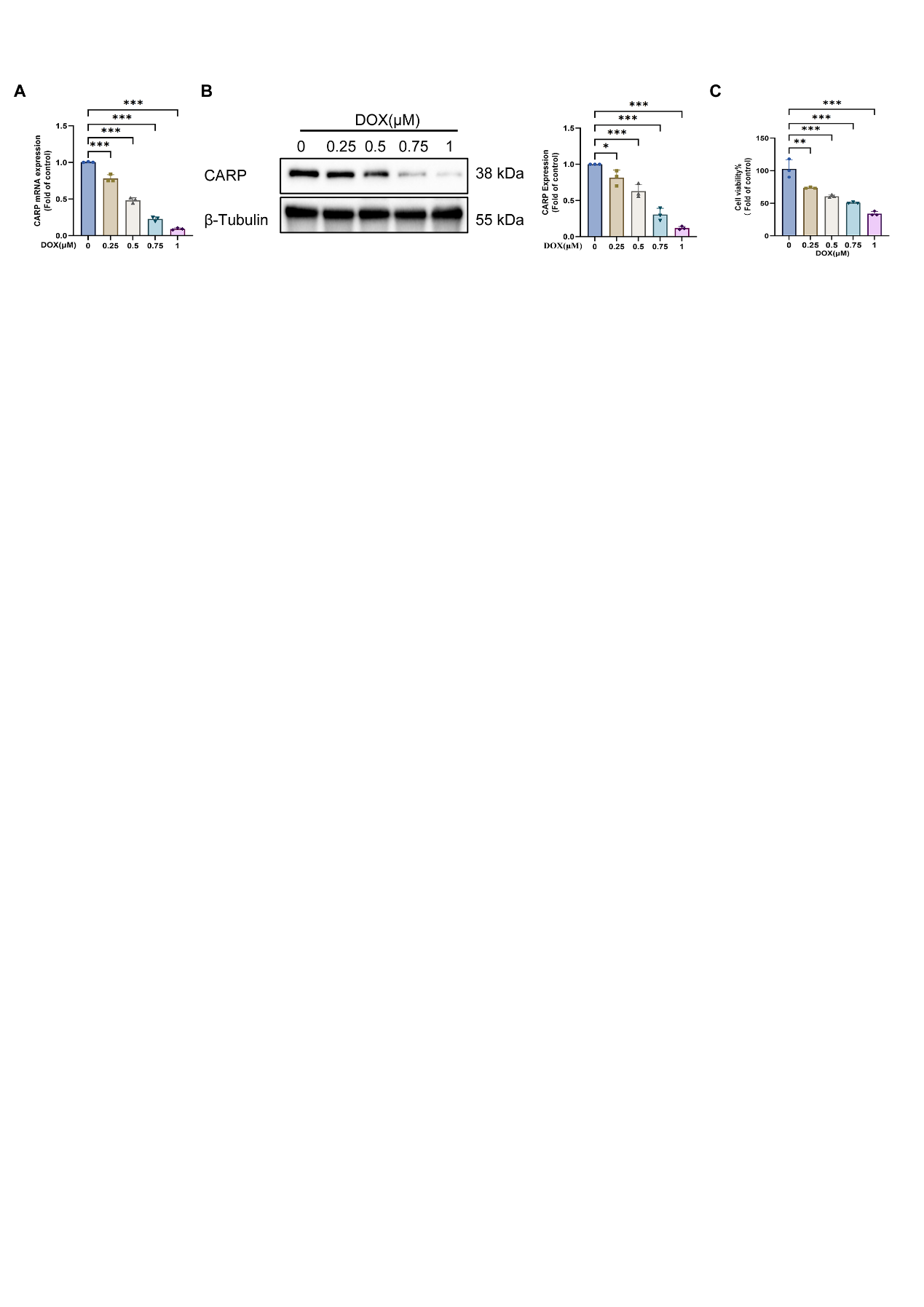

**Fig S4 Dose-dependent inhibition of CARP expression and cell viability by DOX treatment.**

**(A-B)** Relative mRNA expression levels of CARP and representative Western blot images with quantitative analysis in H9C2 cells following 24 h treatment with different concentrations of DOX.(n = 3). **(C)** Assessment of DOX-induced cytotoxicity in H9C2 cells using the CCK-8 assay (n = 3). Data represent mean ± SEM. Statistical significance was determined by one-way ANOVA with Dunnett multiple comparisons test: **p* < 0.05, ** *p* < 0.01, *** *p* < 0.001; ns, not significant (*p* > 0.05).

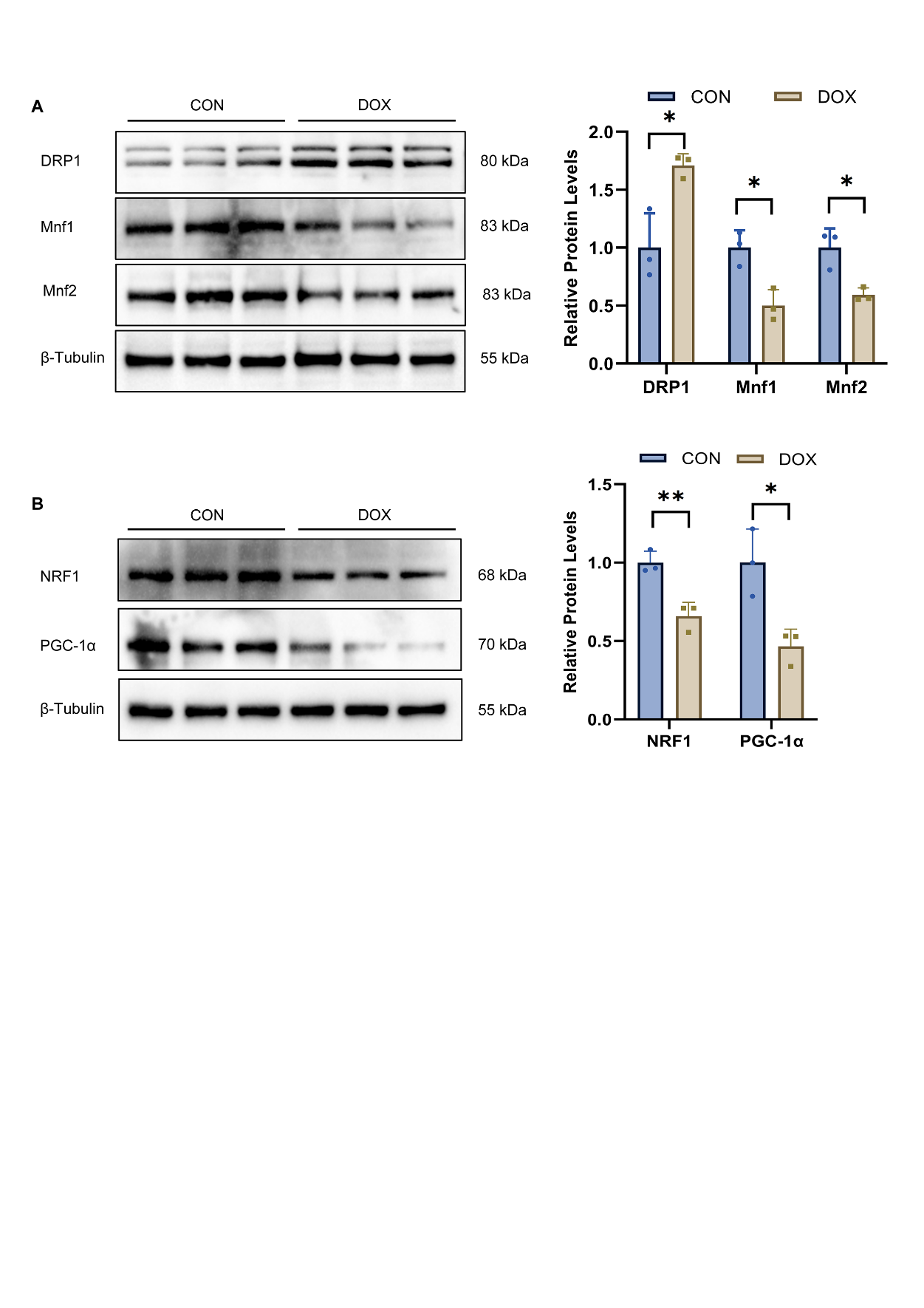

**Fig S5 DOX increases expression of genes involved in promotion of mitochondrial fission while suppressing genes involved in fusion and biogenesis.**

**(A-B)** Western blot analysis of mitochondrial dynamics regulators (DRP1, Mfn1，Mnf2) and biogenesis factors (PGC-1α, NRF1) with corresponding densitometry (n = 3). Data are mean ± SEM. Statistical significance was assessed by two-tailed unpaired t-tests: **p* < 0.05, ** *p* < 0.01, *** *p* < 0.001; ns, not significant (*p* > 0.05).

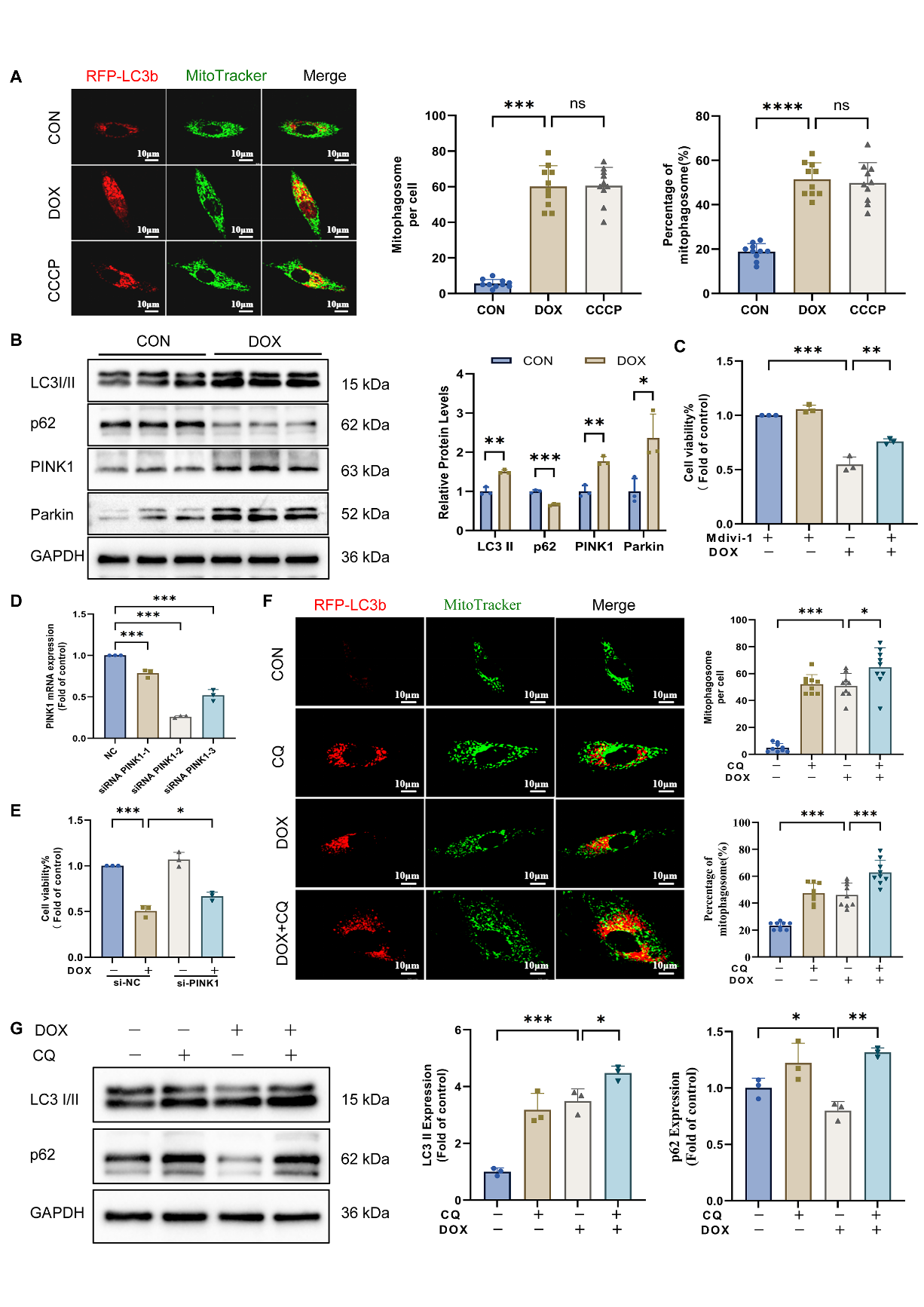

Fig S6 DOX induces mitophagy.

(A) Fluorescent reporter system (RFP-LC3b) labeling autophagosomes. After treatment with 0.75 μM DOX for 24 h followed by 20 μM CCCP for 6 h, MitoTracker Green was used to label mitochondria. Representative confocal microscopy images and quantitative analysis of mitochondrial-autophagosome colocalization are shown, scale bar = 10 μm (n = 10 cells/group). (B) Western blot analysis of mitophagy proteins (LC3-II, P62 PINK1, Parkin) with representative images and quantitation (n = 3). (C) Cell viability was assessed by CCK-8 assay after co-treatment with 0.75 μM DOX and 5 μM Mdivi-1. **(D)** qPCR analysis of PINK1 mRNA levels in H9C2 cells (n = 3). **(E)** Cell viability assessed by CCK-8 assay. **(F)** Fluorescent reporter system (RFP-LC3b) labeling autophagosomes. After treatment with 0.75 μM DOX for 24 h followed by 50 μM CQ for 6 h, MitoTracker Green was used to label mitochondria. Representative confocal microscopy images and quantitative analysis of mitochondrial-autophagosome colocalization are shown, scale bar = 10 μm (n = 10 cells/group). (G) Representative Western blot images and quantitative analysis of LC3-II and P62 protein expression in H9C2 cells treated with 0.75 μM DOX for 24 h followed by 50 μM CQ for 6 h (n = 3). Data are shown as means ± SEM. Statistical significance was assessed by two-tailed unpaired t-tests and one-way ANOVA with Tukey's post hoc test: **p* < 0.05, ** *p* < 0.01, *** *p* < 0.001; ns, not significant (*p* > 0.05).

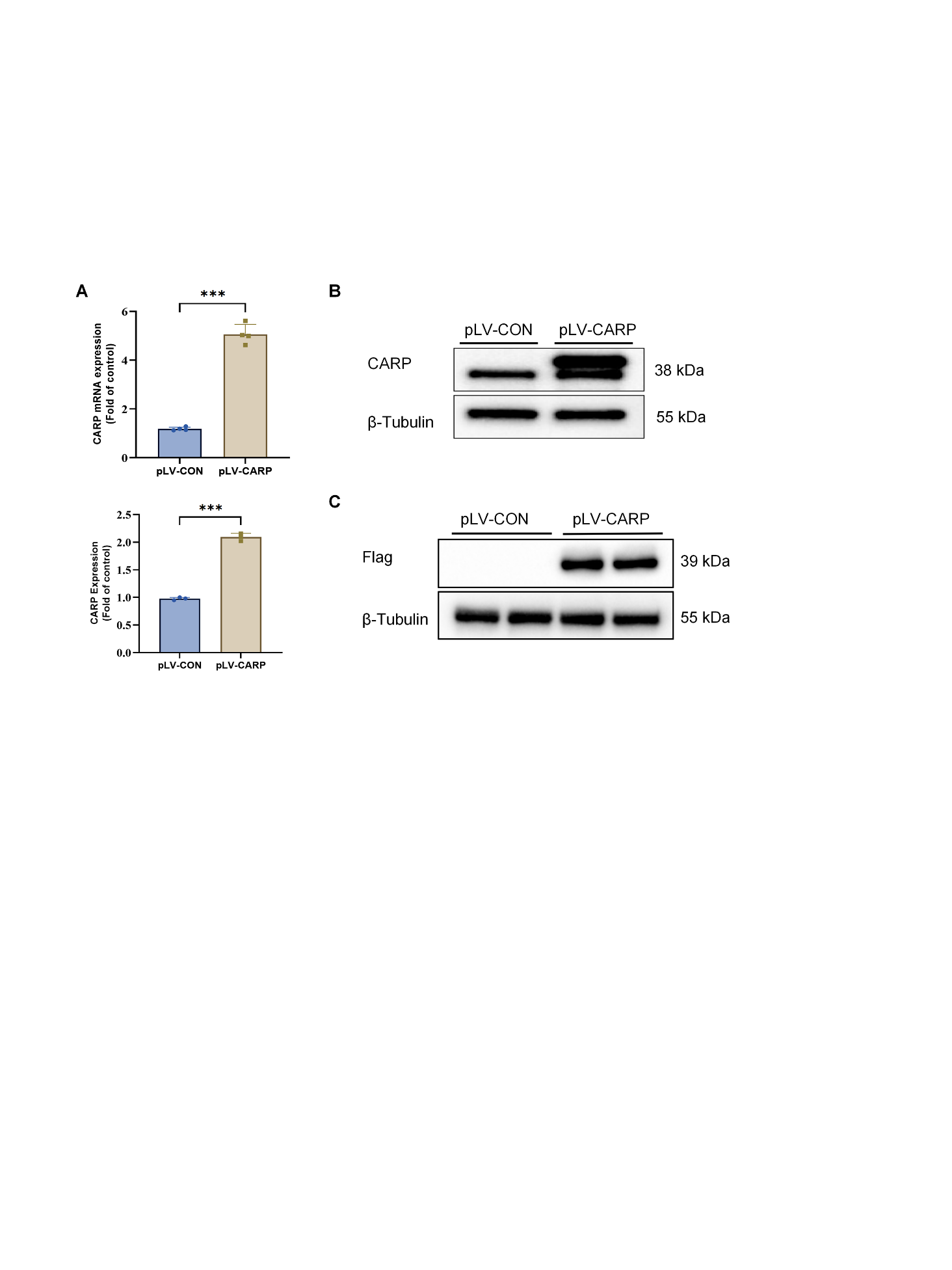

**Fig S7 Establishment of CARP-overexpressing cardiomyocyte line.**

**(A)** qPCR analysis of CARP mRNA levels in H9C2 cells (n = 3). **(B)** Western blot analysis and densitometry of CARP protein expression (n = 3). **(C)** Western blot analysis and densitometry of Flag-tagged protein expression (n = 3). Data are mean ± SEM. Statistical significance was assessed by two-tailed unpaired t-tests: **p* < 0.05, ** *p* < 0.01, *** *p* < 0.001; ns, not significant (*p* > 0.05).

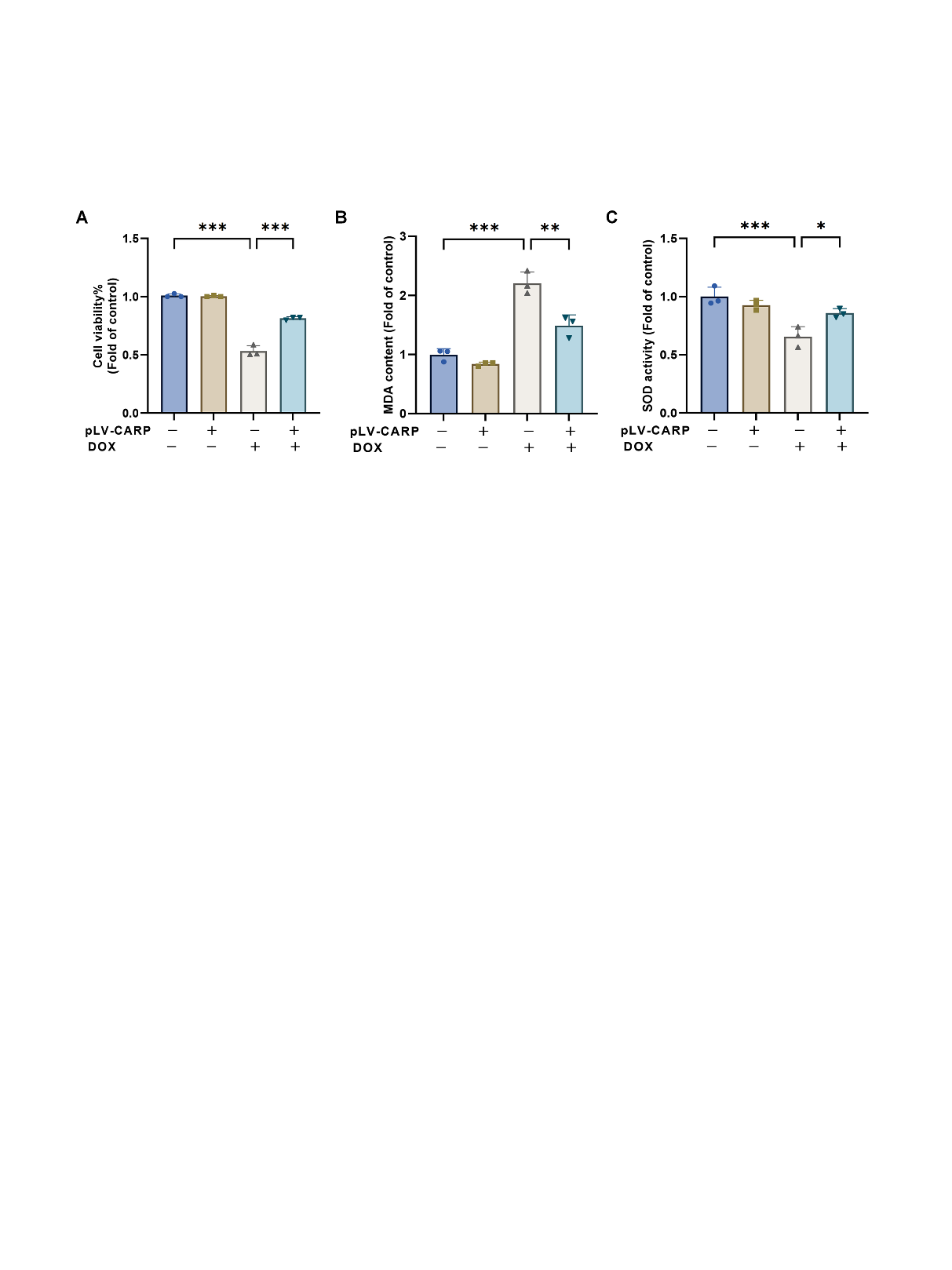

**Fig S8 CARP overexpression attenuates DOX-induced impairment of cardiomyocyte viability and oxidative stress.**

**(A**) Cell viability assessed by CCK-8 assay (n = 3). (B) SOD activity in H9C2 cells (n = 3). (C) MDA content in H9C2 cells (n = 3). Data represent mean ± SEM. Statistical significance was determined by one-way ANOVA with Tukey's post hoc test: **p* < 0.05, ** *p* < 0.01, *** *p* < 0.001; ns, not significant (*p* > 0.05).

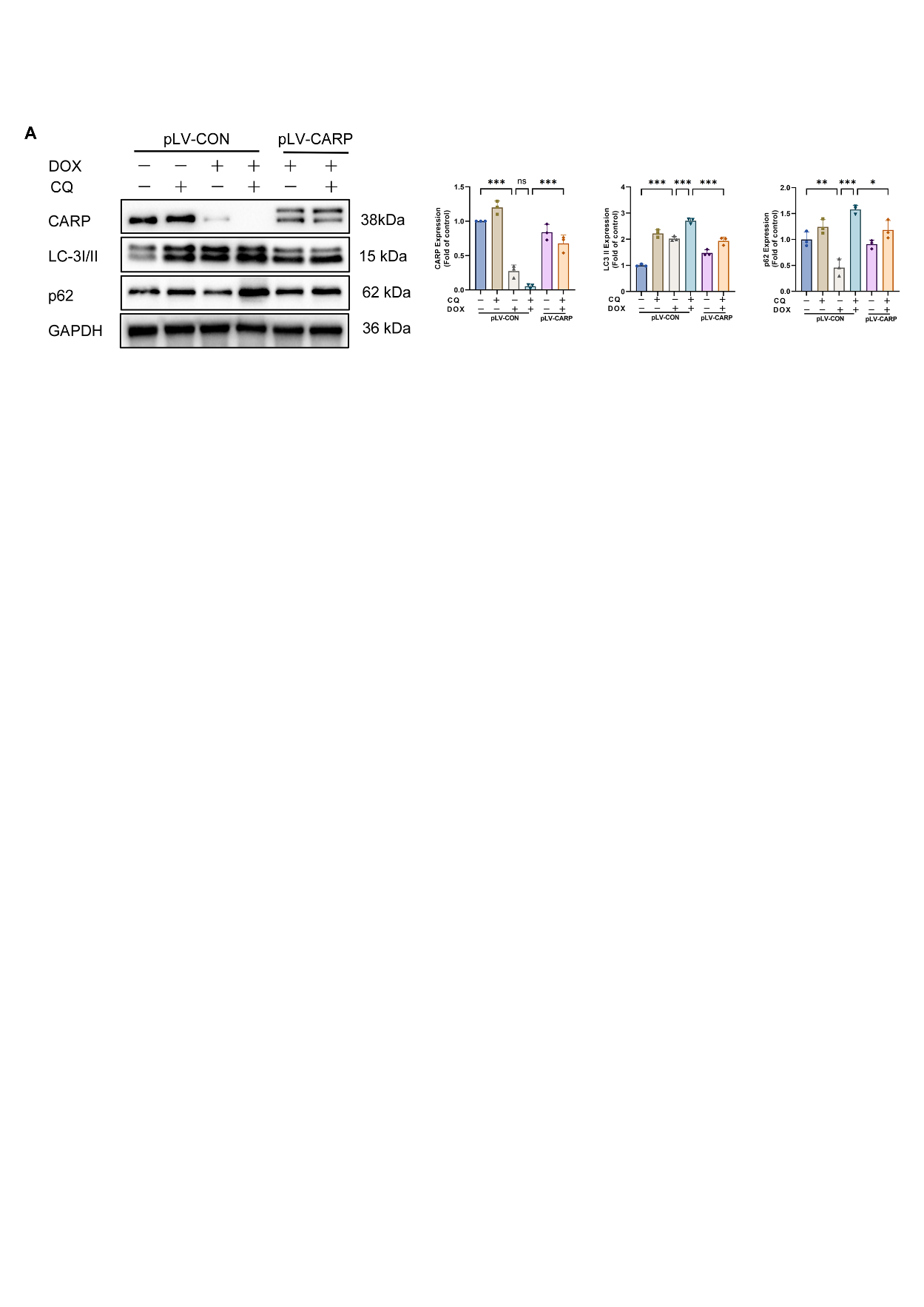

**Fig S9 CARP overexpression inhibits DOX-induced autophagic flux.**

**(A)** Western blot analysis of LC3 II，p62 protein levels with representative images and quantitation (n = 3). Statistical significance was determined by one-way ANOVA with Tukey's post hoc test: **p* < 0.05, ** *p* < 0.01, *** *p* < 0.001; ns, not significant (*p* > 0.05).

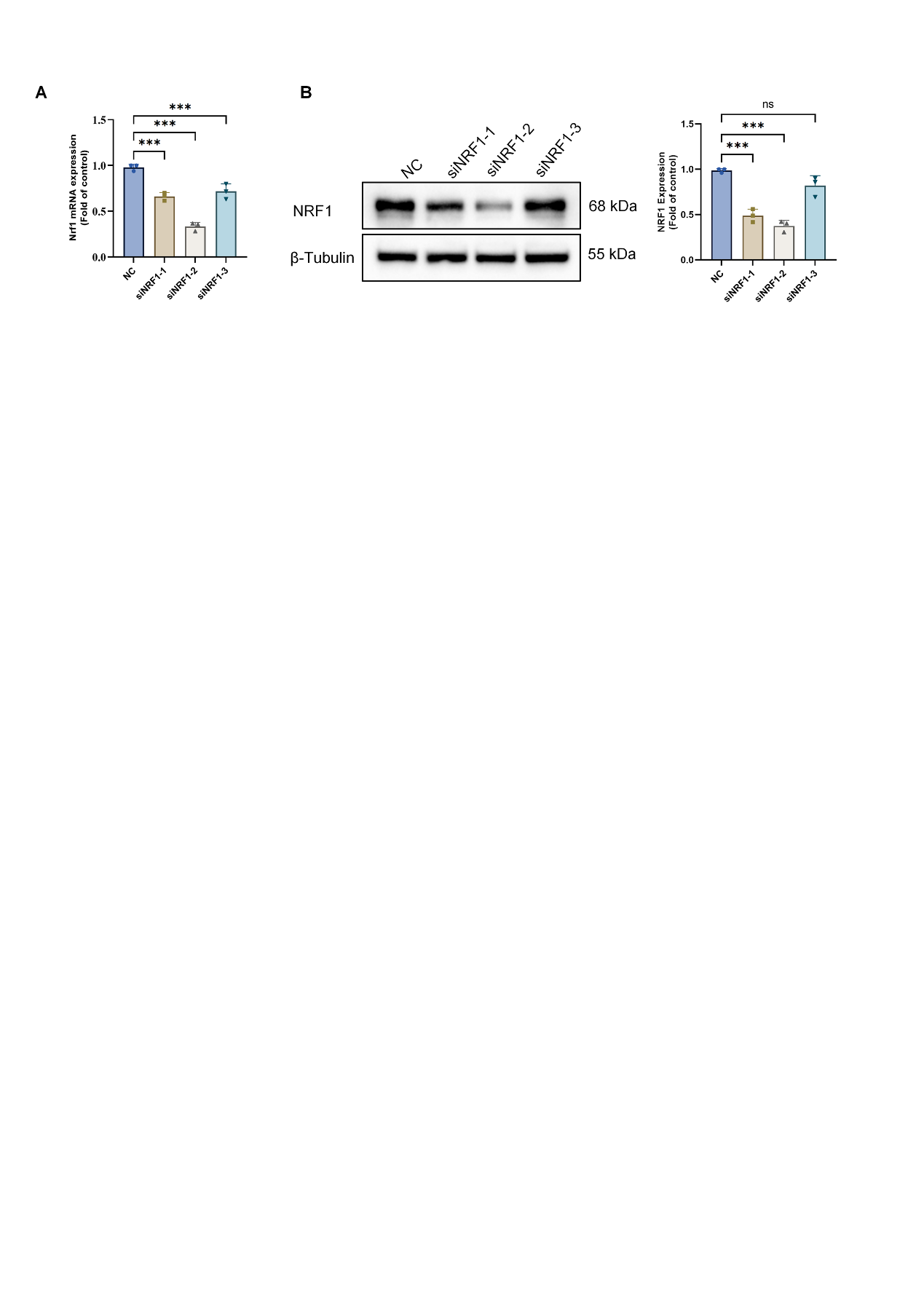

**Fig S10 siRNA-mediated NRF1 knockdown in cardiomyocytes.**

**(A)** qPCR analysis of NRF1 mRNA levels in H9C2 cells (n = 3). **(B)** Western blot analysis and densitometry of NRF1 protein expression (n = 3). Data are mean ± SEM. Statistical significance was determined by one-way ANOVA with Tukey's post hoc test: **p* < 0.05, ** *p* < 0.01, *** *p* < 0.001; ns, not significant (*p* > 0.05).

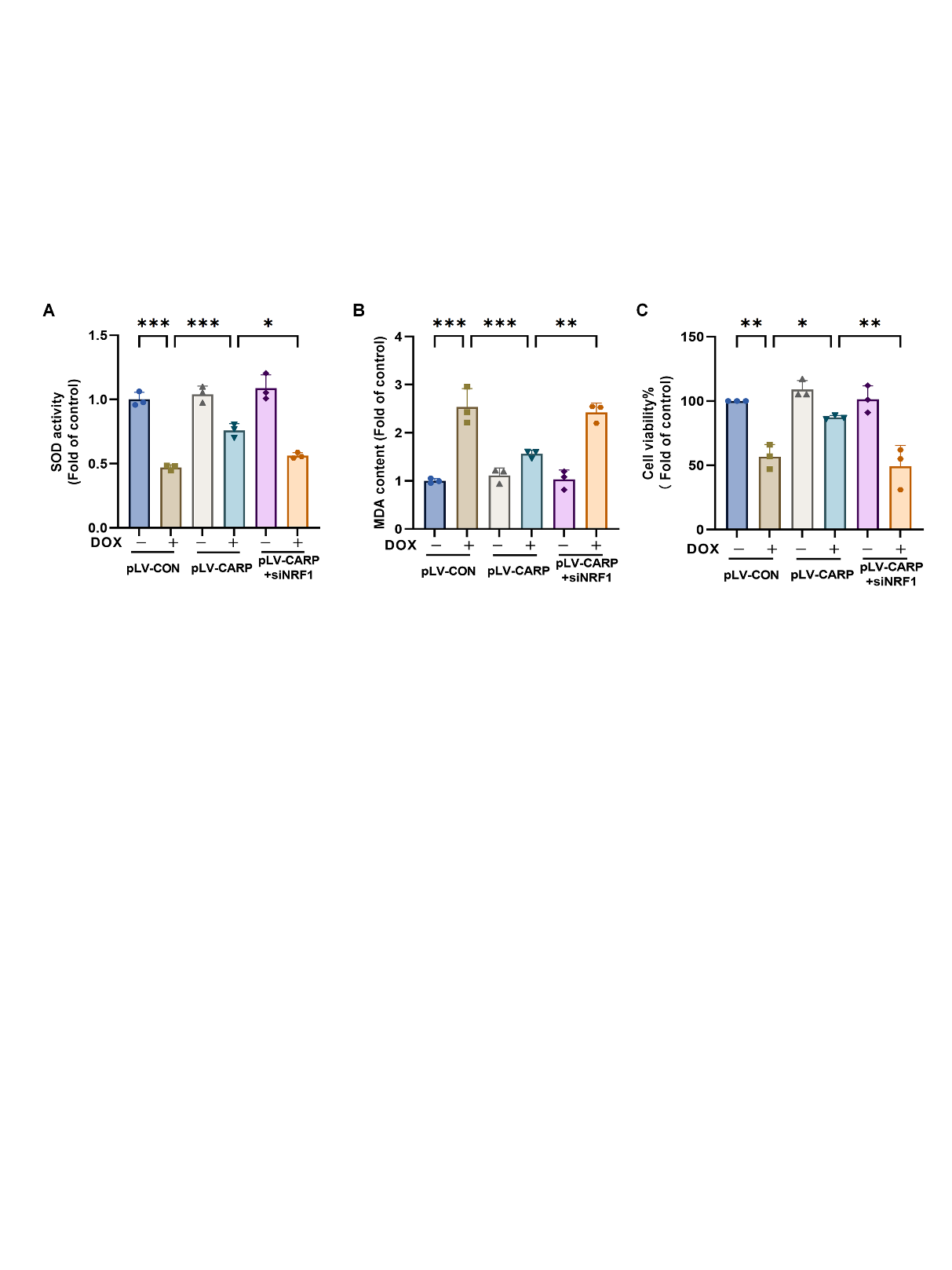

**Fig S11 CARP attenuates DOX-induced impairment of cardiomyocyte viability and oxidative stress via NRF1.**

**(A**) SOD activity in H9C2 cells (n = 3). **(B)** MDA content in H9C2 cells (n = 3).**(C)** Cell viability assessed by CCK-8 assay (n = 3). Data represent mean ± SEM. Statistical significance was determined by one-way ANOVA with Tukey's post hoc test: **p* < 0.05, ** *p* < 0.01, *** *p* < 0.001; ns, not significant (*p* > 0.05).

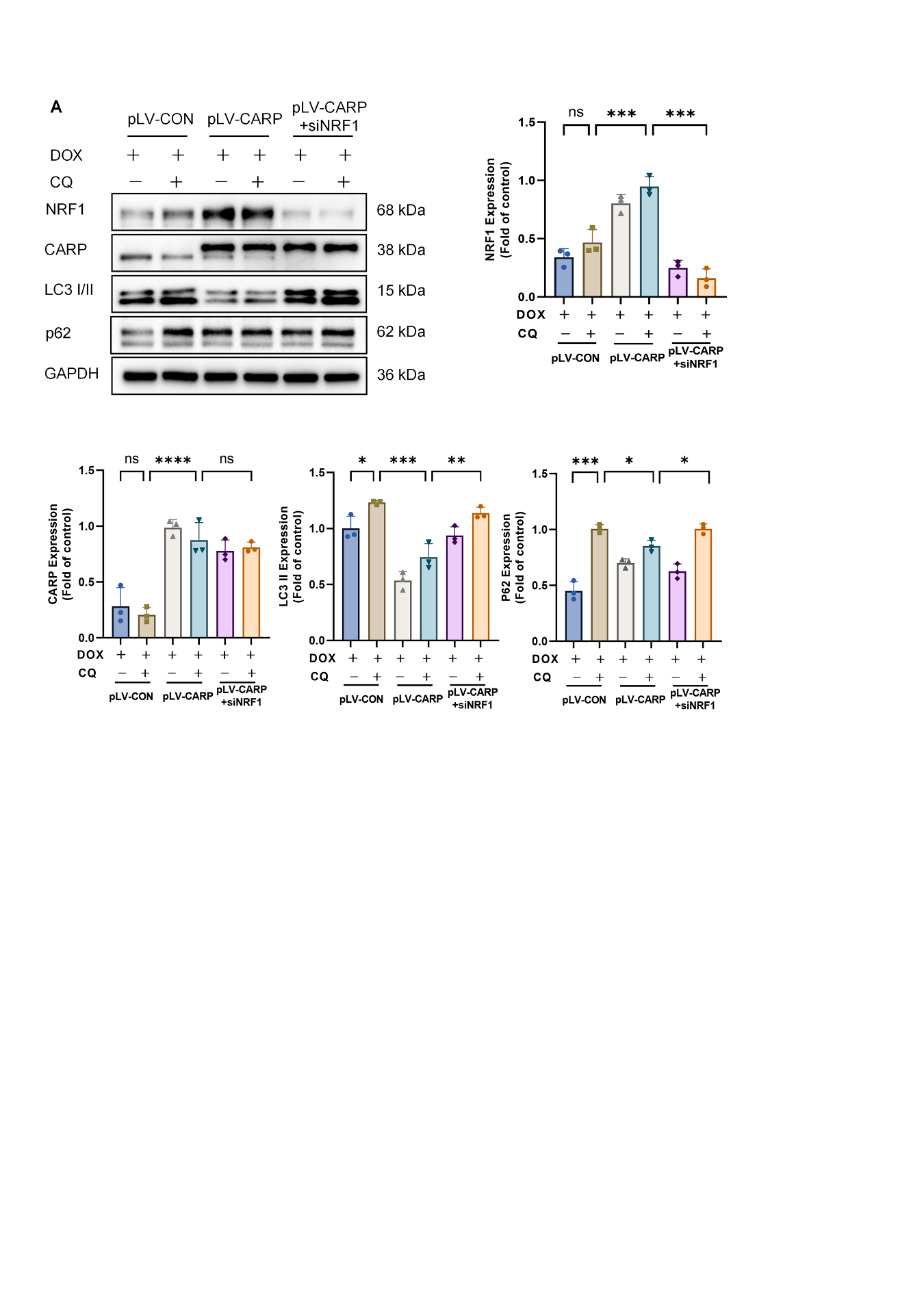

**Fig S12 CARP suppresses DOX-induced autophagic flux through NRF1-dependent mechanisms**.

(A) Western blot analysis of LC3 II，p62 protein levels with representative images and quantitation. Data represent mean ± SEM. Statistical significance was determined by one-way ANOVA with Tukey's post hoc test: **p* < 0.05, ** *p* < 0.01, *** *p* < 0.001; ns, not significant (*p* > 0.05).
